# Mechanisms Underlying the Enhanced Biomass and Abiotic Stress Tolerance Phenotypes of an Arabidopsis MIOX Over-expresser

**DOI:** 10.1101/558239

**Authors:** Nirman Nepal, Jessica P. Yactayo-Chang, Karina Medina-Jiménez, Lucia M. Acosta-Gamboa, María Elena González-Romero, Mario A. Arteaga-Vázquez, Argelia Lorence

## Abstract

*Myo*-inositol oxygenase (MIOX) is the first enzyme in the inositol route to ascorbate (_L_-ascorbic acid, AsA, vitamin C). We have previously shown that Arabidopsis plants constitutively expressing *MIOX* have elevated foliar AsA content and displayed enhanced growth rate, biomass accumulation, and increased tolerance to multiple abiotic stresses. In this work, we used a combination of transcriptomics, chromatography, microscopy, and physiological measurements to gain a deeper understanding of the underlying mechanisms mediating the phenotype of the *At*MIOX4 line. Transcritpomic analysis revealed increased expression of genes involved in auxin synthesis, hydrolysis, transport, and metabolism, which are supported by elevated auxin levels both *in vitro* and *in vivo*, and confirmed by assays demonstrating their effect on epidermal cell elongation in the *At*MIOX4 over-expresser plants. Additionally, we detected up-regulation of transcripts involved in photosynthesis that was validated by increased efficiency of the photosystem II and proton motive force. We also found increased expression of amylase leading to higher intracellular glucose levels. Multiple gene families conferring plants tolerance to cold, water limitation, and heat stresses were found to be elevated in the *At*MIOX4 line. Interestingly, the high AsA plants also displayed up-regulation of transcripts and hormones involved in defense including jasmonates, defensin, glucosinolates, and transcription factors that are known to be important for biotic stress tolerance. These results overall indicate that elevated levels of auxin and glucose, and enhanced photosynthetic efficiency in combination with up-regulation of abiotic stresses response genes underly the higher growth rate and abiotic stresses tolerance phenotype of the *At*MIOX4 over-expressers.

## Introduction

L-Ascorbic acid (AsA) also known as vitamin C is important for human, animal, and plant health. In animals, vitamin C helps in collagen formation, decreases cholesterol content, boosts immune function, and improves wound healing. Vitamin-C deficiency on the other hand leads to scurvy. Humans and other primates are unable to produce vitamin C, so they depend on fruits and vegetables to satisfy their need for this vitamin (Leong 2017; Figueroa-Mendez and Rivas-Arancibia 2015; Weaver et al., 2014). In plants, ascorbate acts as a redox buffer and a cofactor of enzymes involved in various biochemical processes. Ascorbate also participates in vitamin E regeneration and acts as a precursor of organic acids including theronic and tartaric acids (Debolt et al., 2006; Gallie et al., 2013; Gest et al., 2013). Ascorbate modulates the cell cycle, flowering time, and signal transduction affecting plant growth and development (Ortiz-Espín et al., 2018). Vitamin-C deficient (*vtc)* mutants have lower AsA, slower growth, and show early senescence (Conklin et al., 1999; Dowdle et al., 2007; Pastori et al., 2003; Kerchev et al., 2011).

Despite the importance of vitamin C for human health it was until 1998 when a pathway for its biosynthesis in plants was revealed (Wheeler et al., 1998). Two additional routes contributing to the formation of AsA, the L-gulose and D-galacturonate pathways, were proposed in 2003 (Agius et al., 2003; Wolucka and Montagu 2003) and an additional route that uses *myo*-inositol as a precursor was proposed by Lorence et al (2004) (Figure 1).

**Figure 1.**
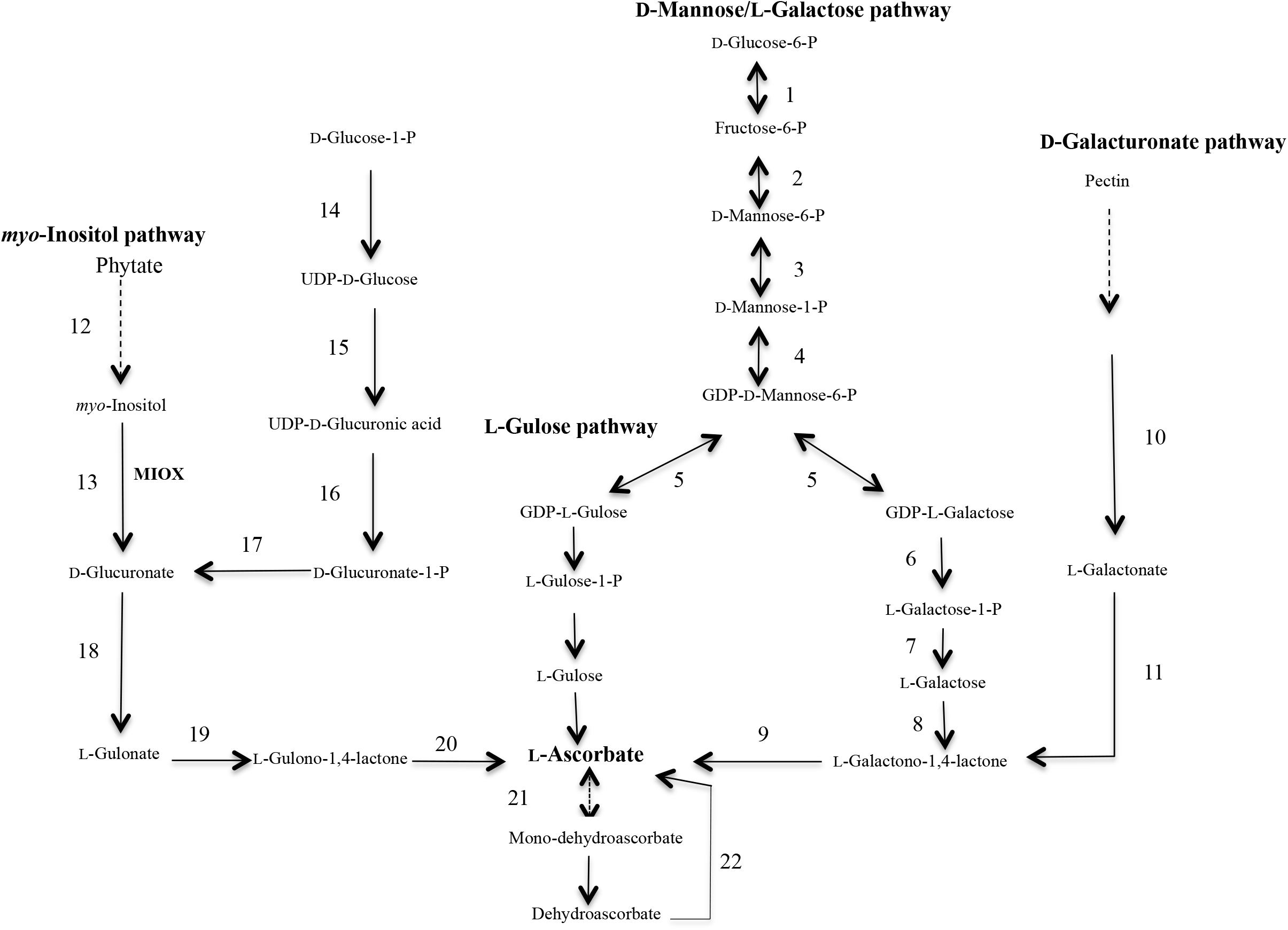
The ascorbate metabolic network in plants and animals. The Mannose/Galactose route, the L-Gulose shunt, the D-Galactouronate route and the *myo*-Inositol pathway. Enzymes: 1 Phosphoglucose isomerase: 2 Phosphomannose isomerase; 3 Phosphomannose mutase 4 GDP-Mannose pyrophosphorylase; 5 GDP-Mannose,3’5’ epimerase; 6 GDP-Galactose phosphorylase,; 7 L-Galactose-1-P-phosphatase; 8 L-Galactose dehydrogenase,; 9 L-Galactono-1,4-lactone dehydrogenase; 10 Galactouronate reductase; 11 Aldonolactonase; 12 *myo*-Inositol 1-P Phosphatase; 13 *myo*-Inositol oxygenase; 14 UDP-Glucose pyrophosphorylase; 15 UDP-Glucose dehydrogenase 16 UDP-Glucuronate pyrophosphorylase;17 Glucuronate 1-kinase; 18 Glucuronate reductase; 19 Glucuronolactonase; 20 L-Gulono-1,4-lactone oxidase; 21 Monodehydroascorbate reductase; 22 Dehydroascorbate reductase.

Ascorbate is the most abundant antioxidant in plants present at concentrations orders of magnitude higher than others. Ascorbate scavenges reactive oxygen species (ROS) and protects plants from abiotic and biotic stresses by maintaining ROS homeostasis (Foyer and Noctor 2011). Plants with elevated AsA content have been developed by over-expressing biosynthetic genes and master regulators (reviewed in Yactayo-Chang et al., 2018). Plants with lower vitamin C (*vtc* mutants) are sensitive to high light, heat, osmotic, and oxidative stresses (Conklin et al., 2013; Cho et al., 2016; Tóth et al., 2011), but are resistant to *Pseudomonas syringae* infection due to increased expression of pathogen-related (PR) genes via reduced glutathione accumulation (Pavet et al., 2006). Contrastingly, enhanced resistance of low ascorbate *A. thaliana* to *P. syringae* is due to higher ROS priming with increased salicylic acid (SA) and enhanced expression of SA induced PR proteins (Mukherjee et al., 2010). Plants with elevated AsA on the other hand are tolerant to salt, cold, heat, osmotic, oxidative, high light and water limitation stresses (reviewed in Yactayo-Chang et al., 2018).

Ascorbate acts as cofactor for the thioglucosidase degradation enzyme, myrosinase that produces toxic chemicals against herbivores and pathogens (Burmeister et al., 2000), and there are reports indicating that AsA can also acts as an antioxidant agent and protect caterpillars from ROS generated from the oxidation of ingested polyphenols (Barbehenn et al., 2002). Other work reported that when the peach potato aphid (*Myzus persicae*) was allowed to feed on potato leaves with normal AsA content, the aphid grew twice the size of its counterpart growing on leaves with low AsA content, indicating that elevated AsA may aid in aphid growth (Kerchev et al., 2013). When Arabidopsis roots were infected with the nematode *Heterodera schachtii*, MIOX played a significant role in the development of syncytium and female nematodes. The study of metabolites in roots containing syncytia showed that, a decrease in the level of *myo*-inositol was responsible for syncytium development (Siddique et al., 2009, Siddique et al., 2014). These examples highlight the species-specific and context-dependent versatile role of AsA during biotic stress.

Arabidopsis plants constitutively expressing enzymes in the *myo*-inositol pathway: *myo*-inositol oxygenase, glucuronate reductase, or L-gulono-1,4-lactone oxidase (*GLOase*, a.k.a. *GulLO*) have 1.5-3 fold elevated AsA content compared to wild type (WT) control (Radzio et al., 2003; Lorence et al., 2004; Tóth et al., 2011; Yactayo-Chang 2011). Using manual phenotyping we and others have showed that Arabidopsis *MIOX4* and *GulLO* over-expressers also display enhanced biomass accumulation of both aerial and underground tissues, and increased tolerance to cold, heat, salt, and pyrene, an environmental pollutant (Lisko et al., 2013; Tóth et al., 2011). Using high throughput phenotyping approaches we have quantified the enhanced growth rate and biomass accumulation of the high AsA line (Lisko et al., 2013; Lisko et al., 2014; Yactayo-Chang et al., 2018). In this study we report our findings on the diverse mechanisms underlying the enhanced growth rate, biomass accumulation, and broad abiotic stress tolerance of the high AsA line using a combination of transcriptomics (RNA-seq), RT-qPCR, liquid chromatography-tandem mass spectroscopy (LC-MS/MS), cell biology, confocal microscopy, and physiological measurements (Figure 2).

**Figure 2.**
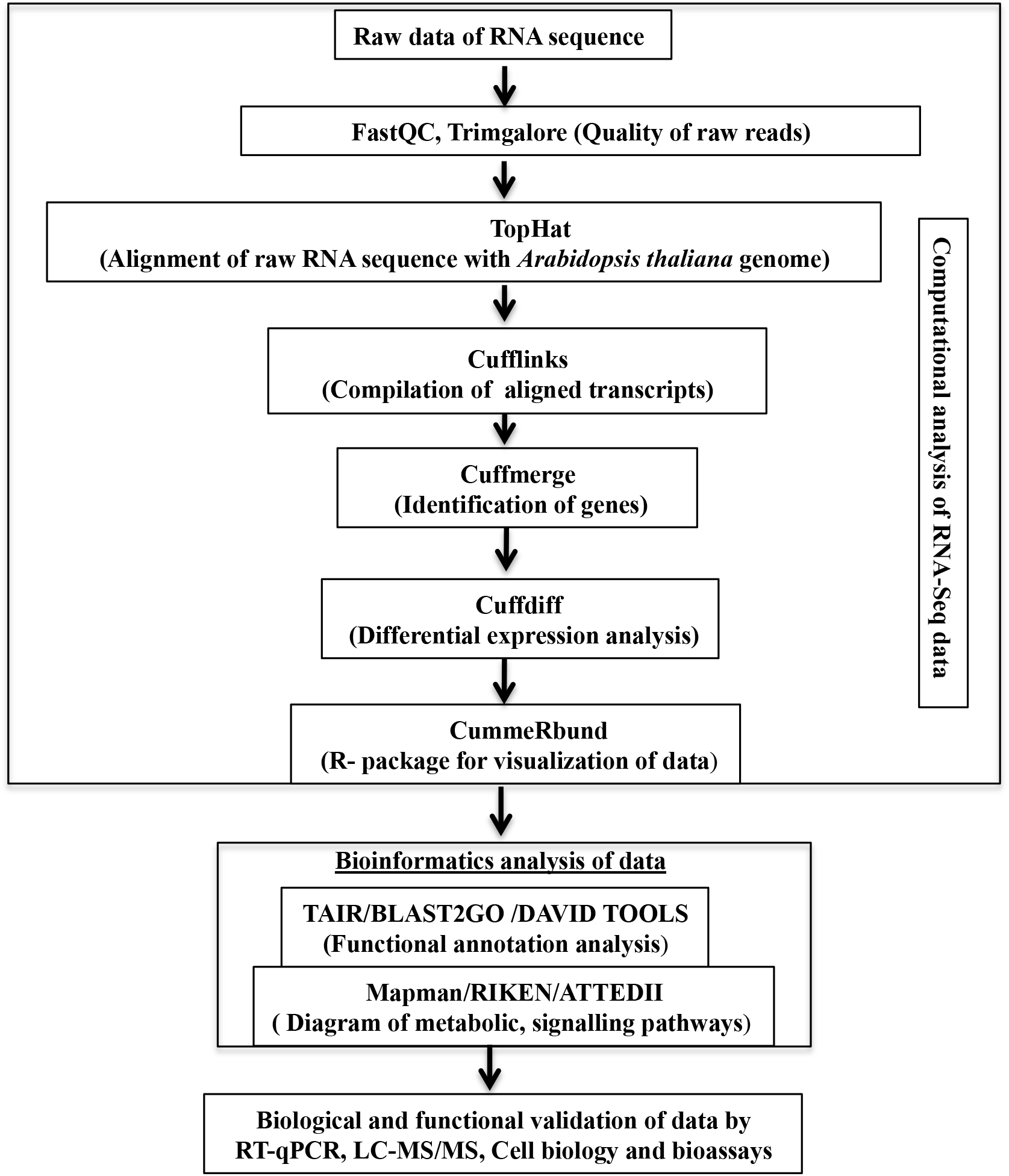
Schematic representation of RNA-Seq data analysis. Raw RNA Sequence data was analyzed by using TopHat, Cufflink, Cuffmerge, Cuffdiff and CummeRbund algorhitms. Functional annotation and metabolic pathways of transcriptomics datasets were analyzed by BLAST2gOv3.1 and Mapman. RT-qPCR validation of hallmark genes expression, quantification of metabolites by LC-MS/MS and completion of data analysis.

## Materials and methods

### Plants and growth conditions

In this study, we used *Arabidopsis thaliana (A. thaliana)* wild type accession Columbia (ABRC stock CS-60000), single-insertion homozygous line over-expressing *At*MIOX4 (*At*MIOX4 L21, Yactayo-Chang 2011), a vitamin-C defective mutant (*vtc1-1*) (Conklin et al., 1999) and the auxin sensor line (R2D2) (Liao et al., 2015). Seeds were sterilized with 70% (v/v) ethanol for 5 min followed by 50% (v/v) sodium hypochlorite with 0.05% (v/v) Tween-20 for 10 min. Then, seeds were washed 5-6 times with sterile distilled water. Plants were grown in Murashige and Skoog (MS) media pH 5.6 for 12 days after stratification for 3 days at 4°C. Seedlings from MS plates were transferred to soil (Promix PGX; Premier Horticulture) and grown in a growth chamber at 23°C, 65% humidity, 165-210 μmol/m^2^/s light intensity and 10:14 h photoperiod.

### Southern blot

Genomic DNA of the *At*MIOX4 OE line was isolated using the CTAB method (Murray and Thompson 1980). Briefly, 4-5 g of fresh tissue was grinded in a mortar and pestle cooled with liquid nitrogen. Then, 20 ml of CTAB buffer (1 M Tris-HCl pH 8.0, 5 M NaCl, 0.5 M EDTA, 2% CTAB, and 1% (v/v) β-mercaptoethanol) were added, vortexed and incubated at 65°C for 20 min. After incubation, 20 ml of chloroform: isoamyl alcohol 24:1 (v/v) were added. The lysates were gently mixed and centrifuged at 4,000xg at 4°C for 20 min. The upper aqueous phase was transferred to new tube and, total DNA was extracted and precipitated with 3M NaAc pH 5.2. The precipitated DNA was dissolved in TE buffer. Five micrograms of extracted DNA were digested with *At*MIOX4:pBIB-Kan with *Sac*I or *Kpn*I. Digested DNA was separated on 0.8% (w/v) agarose 1X TAE gel and electro blotted onto a positively charged nylon membrane (Roche), using 1X TBE as transfer buffer. The UV-cross-linked membrane was then pre-hybridized at 65°C for 15 min in Church buffer (sodium phosphate buffer containing, 0.5 M Na_2_HPO_4_ pH 7.2, 20% (w/v) SDS, 1 mM EDTA, and 1% BSA), and hybridized overnight at 65°C with 50 ng of DNA probe each labeled with 50 *μC*i[^32^P]-dCTP. The probes were prepared using Prime-It^®^ Rm T Random Primer Labeling kit (Stratagene). The probe for *At*MIOX4:pBIB-Kan was amplified with npt-II forward (5’-AGA GGC TAT TCG GCT ATG AC-3’) and npt-II reverse (5’-AGC TCT TCA GCA ATA TCA CG-3’) primers at 55°C annealing temperature. After hybridization, the blots were washed with phosphate buffer (0.5 M Na_2_HPO_4_, 20% (w/v) SDS, 1 mM EDTA, and 33 mM NaCl pH 7.2 at 65 °C. Photographs were taken by exposing membranes to film for 2-3 days at −80°C (Biomax XAR, Kodak).

### RNA extraction and sequencing

Three biological replicates of leaf samples were collected when plants completed the vegetative growth stage (stage 5.0 according to Boyes et al., 2001). Total RNA was purified using the RNeasy kit (Qiagen) following the manufacturer’s instructions. RNA quality and quantity were assessed with an Experion system using the Experion™ RNA HighSens chips (Bio-Rad). RNA samples were submitted to the Core Facility at Michigan State University for sequencing. Six libraries were prepared and sequenced using manufacturer recommendations with a Hi-Seq Illumina 2500 system. Libraries were prepared using the Illumina TruSeq Stranded mRNA Library Preparation Kit. After QC and quantitation the libraries were combined into a single pool for multiplexed sequencing. The pool was loaded on two lanes of an Illumina HiSeq 2500 Rapid Run flow cell (v2). Sequencing was performed in a 1×50bp single read format. Base calling was done by Illumina Real Time Analysis (RTA) v1.18.64 and output of RTA was de-multiplexed and converted to FastQ format with Illumina Bcl2fastq v1.8.4. Sequencing data used in this study (#SRA accession:PRJNA515221) will be available upon publication.

### Bioinformatics analysis of raw RNA-Seq data

Quality analysis of raw data was performed using FastQC v0.11.7 (Andrews 2014). Trim Galore v0.4.3.1 was used for trimming the Illumina universal adapter sequences and removal of reads less than 40 bp in length (Krueger 2015). TopHat v 2.1.1 was used for aligning the reads with the whole genome sequence of *A. thaliana* genome (TAIR10) (https://www.arabidopsis.org/) using maximum edit distance 2.0, library type unstranded, maximum intron length 10,000 bp, minimum intron length 50 bp and default parameters (Kim et al., 2013). Cufflinks (v2.2.1) and Cuffmerge (v3.0) algorithms were used next for transcript compilation and gene identification using *A. thaliana* gene annotation TAIR10_gff3/TAIR10_GFF3_genes_transposons.gff (https://www.arabidopsis.org/). Differential expression analysis was performed using the CuffDiff (v7.0) algorithm. The CummeRbund algorithm R package (v2.22.0) was used for data visualization (Trapnell et al., 2012). Differentially expressed (DE) transcripts were selected based on an absolute value of log2 ratio ≥1.5 with FDR adjusted p-value ≤ 0.05). False discovery rate correction (FDR) was performed using the Benjamini Hochberg method. Blast2GO (Conesa and Gőtz, 2008) and DAVID v6.8 (Huang et al., 2009) were used for gene ontology and functional enrichment analysis with a false discovery rate correction using the Benjamini-Hochberg method with FDR adjusted p-value ≤ 0.05. MapMan 3.5.1R2 (Usadel et al., 2012Vitamin-C deficient) was employed for visualizing up-and down-regulated genes affecting metabolism in plants mapping with Ath_AGI_LOCUS_TAIR10_Aug2012 (https://www.arabidopsis.org/).

### Real-time quantitative PCR (RT-qPCR) for gene expression analysis

Real-time quantitative PCR (RT-qPCR) was used for biological validation of the RNA-seq data using reference genes previously published and following MIQE guideline (Czechowski et al., 2005; Bustin et al., 2008). Three biological replicates of leaf samples were collected when plants completed the vegetative growth stage. Total RNA was purified using the Purelink™ RNA mini kit (Ambion, Life Technologies) following the manufacturer’s instructions. The RNA quality and quantity were assessed with an Experion system using the Experion™ RNA HighSens chips (Bio-Rad). Residual DNA in the sample was removed using DNA-*free*™ DNA removal kit (Invitrogen). Complimentary DNA (cDNA) were prepared by following manufacturer’s instruction (iScript select cDNA synthesis kit, Bio-Rad). Validated primers were obtained from *At*RTPrimer database (Han and Kim 2006). The primers for photosystem II (PSII) related center transcripts and glutathione reductase (*GRX480*) were validated using the guidelines by Bustin et al., 2008. RT-qPCR was performed following manufacturer’s instruction (SsoFast™ EvaGreen^®^ supermixes, Bio-Rad) in a CFX384TM real-time system (Bio-Rad). Normalization of transcripts counts was obtained using ACTIN2 and UBQ10 as internal controls. The relative expression of the genes was calculated using 2^−δδt^ method (Livak and Schmittgen 2001). Three biological replicates and two technical replicates were used for RT-qPCR. Two-way student’s t-test was performed at 0.05 significance level for normalized expression value.

### Di-nitro salicylic acid (DNS) assay for reducing sugars estimation

Six biological replicates of leaf samples were collected when plants completed the vegetative growth stage. Reducing sugars were estimated using di-nitro salicylic assay (DNS). Plant samples were ground and extracted twice in 1 ml of 0.5 mM sodium acetate buffer (pH 5.0) using Precellys 24 lyzer (Bertin Technologies) and 100 μl of DNS reagent (Sigma) was added and incubated in boiling water-bath (95°C) for 20 min. The reaction was stopped by adding 300 μl ddH_2_O. The 125 μl of reaction mixture was transferred into a 96-well clear plastic reading plate with 2 technical replicates and absorbance was measured at 540 nm in a 96 well plate reader (BioTek). Two-way student’s t-test was performed at a 0.05 significance level. The experiment was replicated twice with similar results.

### Glucose oxidase-glucose peroxidase (GOD-POD) assay for estimation of glucose

Five biological replicates of leaf samples were collected when plants completed the vegetative growth stage. Plant samples were ground and extracted twice in 1 ml of 0.5 mM sodium acetate buffer pH 4.5 using Precellys 24 lyzer (Bertin Technologies). Sixty microliters of sample, 54 μl of GOD-POD enzyme mix (Sigma)) and 6 μl of O-dianisidine were mixed and incubated at 37°C for 30 min. The reaction was stopped by adding 60 μl of 10 M HCl. Absorbance was recorded at 540 nm in a 96-well clear plastic reading plate with two technical replicates in a 96 well plate reader (BioTek). Two-way student’s t-test was performed at 0.05 significance level. The experiments were replicated twice with similar results.

### Photosynthetic efficiency measurements

Plants were grown on MS media in 23°C, 65% humidity, 165-210 μmol/m^2^/s light intensity and 10:14 h photoperiod after stratification for 3 days. Twelve days after germination plants were transferred to Arabidopsis soil (Promix PGX; Premier Horticulture LTEE). MultispeQ v1.0 (Kuhlgert et al., 2016) was used to measure the photosystem II efficiency (ΦII) and proton motive force (vH^+^) 21, 23, 26 and 28 days after germination. Fifteen biological replicates for each genotype were measured. Experiment was replicated twice with similar results. Two-way student’s t-test was performed at a 0.05 significance level for each day of measurement.

### Hormones and metabolites quantification using LC-MS/MS

Leaf samples from 10 biological replicates for each genotype were collected and lyophilized in liquid nitrogen when plants completed the vegetative growth stage. Hormones extraction and analysis were performed at the University of Nebraska, Lincoln with modifications of Lan et al., 2014 and Westfall et al., 2016. Briefly, lyophilized samples were ground using a Tissuelyzer (Qiagen). Samples were spiked with deuterium–labelled internal standards and extracted in ice-cold methanol:acetonitrile (MeOH:CAN,1:1,v/v) for 2 min at 15 Hzs^−1^ then centrifuged at 16,000xg for 5 min at 4°C. Supernatants were pooled and evaporated in a Labconco Speedvac. The pellets after evaporation were redissolved in 200 μl of 30% MeOH. For LC separation, a ZORBAX Eclipse Plus C18 column (2.1 mm × 100 mm, Agilent) was used flowing at 0.45 ml/min. The gradient of the mobile phases A (0.1% acetic acid) and B (0.1% acetic acid/90% acetonitrile) was as follow: 5% B for 1 min, to 60% B in 4 min, to 100% B in 2 min, hold at 100% B for 3 min, to 5% B in 0.5 min. The LC system was interfaced with a Sciex QTRAP 6500+ mass spectrometer equipped with a TurboIonSpray (TIS) electrospray ion source. Analyst software (version 1.6.3) was used to control sample acquisition and data analysis. The QTRAP 6500+ mass spectrometer was tuned and calibrated according to the manufacturer’s recommendations. All hormones were detected using MRM transitions that were previously optimized using standards. The instrument was set-up to acquire in positive and negative ion switching. For quantification, an external standard curve was prepared using a series of standard samples containing different concentrations of unlabeled hormones and fixed concentrations of deuterium-labeled standards (D5-IAA, D5-tZ, D5-tZR, D2-JA, D6-ABA, D4-SA). Two-way student’s t-test was performed at 0.05 significance level.

### Semi-quantification of in-vivo auxin

Crosses between *At*MIOX4 L21 (♂) and auxin sensor (R2D2) (♀) were used for visualization and *in-vivo* semi-quantitative analysis of auxin. Plants were grown on MS media in 23°C, 65% humidity, 165-210 μmol/m^2^/s light intensity and 10:14 h photoperiod after stratification for 3 days. Twelve days after germination plants were transferred to Arabidopsis soil (Promix PGX; Premier Horticulture). *At*MIOX4 L21 and R2D2 line were crossed and F1 generation were selected using kanamycin as a selectable marker and PCR using primers for MIOX4 and kanamycin resistance genes. The positive crosses were taken for the *in-vivo* auxin quantification. The positive crosses were grown on MS media horizontally and vertically for *in-vivo* auxin quantification of shoots and roots respectively. The plants were grown at 23°C, 65% humidity, 165-210 μmol/m^2^/s light intensity and 10:14 h photoperiod for 7 days. The seedlings were mounted in propidium iodide (10 μg/ml) for root scanning and in PBS for shoots scanning. The image acquisition and analysis of roots and shots were performed in Cytation™ 5 cell imaging multimode reader and Gen3.03 software (BioTek). Venus was excited at 488 nm and detected at 498-530 nm, ntdTomato was excited at 561 nm and detected at 571-630, propidium iodide was excited at 561 nm and detected at 571-700 nm (Liao et al., 2015).

### Root length, hypocotyl length and hypocotyl cell length measurement

Plants were grown on plates positioned vertically on MS media in 23°C, 65% humidity, 165-210 μmol/m^2^/s light intensity and 10:14 h photoperiod after stratification for 3 days. For primary root length measurement plants were grown on 150-210 μmol/m^2^/s light intensity whereas for hypocotyl measurement plants were grown on dark condition in same growth chamber. Nine days after germination the primary root length and the hypocotyl length were measured with a caliper. Fifteen biological replicates for each genotype were used in this experiment and a two way student’s t-test was performed at 0.05 significance level.

For cell length measurements, hypocotyl of plants grown vertically on MS media on dark conditions were mounted in 1X PBS. The image acquisition and analysis was performed using Cytation™ 5 cell imaging multimode reader (BioTek) and cell length was measured using the Gen3.03 software (BioTek). Hypocotyl epidermal cell length was the average of 6 biological replicates for each genotype. The length of five cells was measured from each sample. Two-way student’s t-test was performed at a 0.05 significance level (Das et al., 2016).

### Ascorbate measurements

Foliar tissue samples (40-60 mg) were collected at developmental stage 5.0 (Boyes et al. 2001) and immediately frozen in liquid nitrogen. The samples were grinded in a cold mortar and pestle using liquid nitrogen and adding 0.75 ml of fresh meta-phosphoric acid (6%) to the sample. The plant extracts were immediately transferred to a 1.5 ml amber tube and centrifuged at 14,000xg for 15 min at 4°C. For the measurement of reduced ascorbate, the supernatants were transferred to 1.5 ml amber tubes and kept on ice. There 0.95 ml of the meta-phosphoric acid buffer was added to a quartz cuvette and 50 μl of supernatant were added and mixed by inverting 2-3 times. The absorbance was recorded at 265 nm. Twenty μl of ascorbate oxidase were added, mixed, and absorbance was recorded at 265 nm. For the measurement of oxidized ascorbate, 0.95 ml of the meta-phosphoric buffer was added into a quartz cuvette and 50 μl of supernatant were added and mixed and absorbance was recorded at 265 nm. The mix was recovered in a 1.5ml amber tube and 1 μl DTT (200 mM) was added mixed and incubated at dark for 20minutes. The final absorbance was recorded at 260 nm. The reduced and oxidized ascorbate content was calculated using the extinction coefficient as [(Total change in absorbance)/14.3]*20*0.75(1/sample weights in grams). The total ascorbate content was calculated as the sum of reduced and oxidized ascorbate pools.

### Quantification of H_*2*_*O*_*2*_ *accumulation*

Foliar tissue samples (40-60 mg) were collected at developmental stage 5.0 and immediately frozen in liquid nitrogen. H_2_O_2_ was quantified using Amplex Red (Molecular Probes, Invitrogen) as previously described (Katano et al. 2017; Suzuki et al. 2015). Briefly, the plant samples were grinded using liquid nitrogen. The Amplex red reagent (50 mM sodium phosphate buffer pH 7.4, 50 μM Amplex red and 0.05 U/ml horse radish peroxidase) was added to grinded tissues and centrifuged at 13,000xg for 12 min at 4°C. After centrifugation, 450 μl of supernatant were transferred to a 1.5ml amber tube and incubated for 30 min in the dark at room temperature. The absorbance was recorded at 560 nm. The concentration of H_2_O_2_ was calculated using a standard curve obtained from 0, 0.5, 1, 3, 6 and 9 μM H_2_O_2_.

### Methione-derived aliphatic glucosinolate HPLC analysis

When plants completed the vegetative growth stage, leaf samples (∼100 mg of DW) from ten biological replicates per genotype were collected and frozen in liquid nitrogen. Glucosinolates extraction was carried out with 80% MeOH and aryl sulfatase (Sigma). Recovered desulfoglucosinolates were lyophilized in Speedvac and redissolved in 500 μl of ultrapure water. Samples were analyzed in a reverse phase HPLC system (Dionex; Ultimate 3000) equipped with an AcclaimTM 300 C18 column (4.6 × 150 mm, 3 μm particle size, 300A°). The mobile phase was composed of acetonitrile (A) and 0.1% TFA in water (B). The gradient program was: 0–1 min, 1.5 % B; 1–6 min, 1.5–5 % B; 6– 8 min, 5–7 % B; 8–18 min, 7–21 % B; 18–23 min, 21–29 % B; 23–23.1 min, 29–100 % B; 23.1–24 min 100 % B, and 24.1–28 min 1.5 % B; flow 1.0 ml/min. Glucoerucin (Nacalai Tesque), 4-methylthiobutil glucosinolate (4MTB), a methionine-derived aliphatic glucosinolate, was analyzed. Detection was performed at 229 nm using a photodiode array. GlucoerucinUV spectra acquisition and analysis were carried following methods described by Grosser and Dam 2017 and Madsen et al., 2016.

### Pathogen infection assay

High ascorbate (*At*MIOX4 L21), low ascorbate (*vtc2-1*) and WT (CS-60000) *A. thaliana* plants were challenged with the virulent pathogen *Pst* DC3000 (*Pseudomonas syringae* pv tomato DC3000). *Pst* DC3000 was grown on King’s B medium for 2 to 3 days and resuspended on MgCl_2_ (10 mM) and Silwet L-77 (0.02%) in 10^5^ colony forming unit (cfu)/ml. Two-week old seedlings grown on MS media were dip inoculated with bacteria and covered for 1 h and uncovered slowly. The amount of bacteria present in plants was analyzed 1 h after dipping (Day 0) and 3 d after dipping (Day 3). Three biological replicates were pooled and measured for bacterial growth as cfu for each genotype (Zhu et al., 2013). The experiment was replicated three times with similar results.

## Results

After years of work we have detected gene silencing in the *At*MIOX4 lines we initially published (Lorence et al., 2004). To continue studying the possible roles of constitutive expression of *At*MIOX4 in plants we decided to develop new transgenic lines. The line used in this study was one of the new ones we developed. *At*MIOX4 line 21 was chosen for this study because it is homozygous, harbors a single copy of the *MIOX4* ORF and recapitulates the phenotype of the lines previously reported (Lorence et al., 2004) that includes elevated levels of AsA (Figure 3C), enhanced biomass and tolerance to multiple abiotic stresses relative to the wild type control (Lisko et al., 2014; Yactayo-Chang, 2011; Yactayo-Chang et al., 2018).

**Figure 3.**
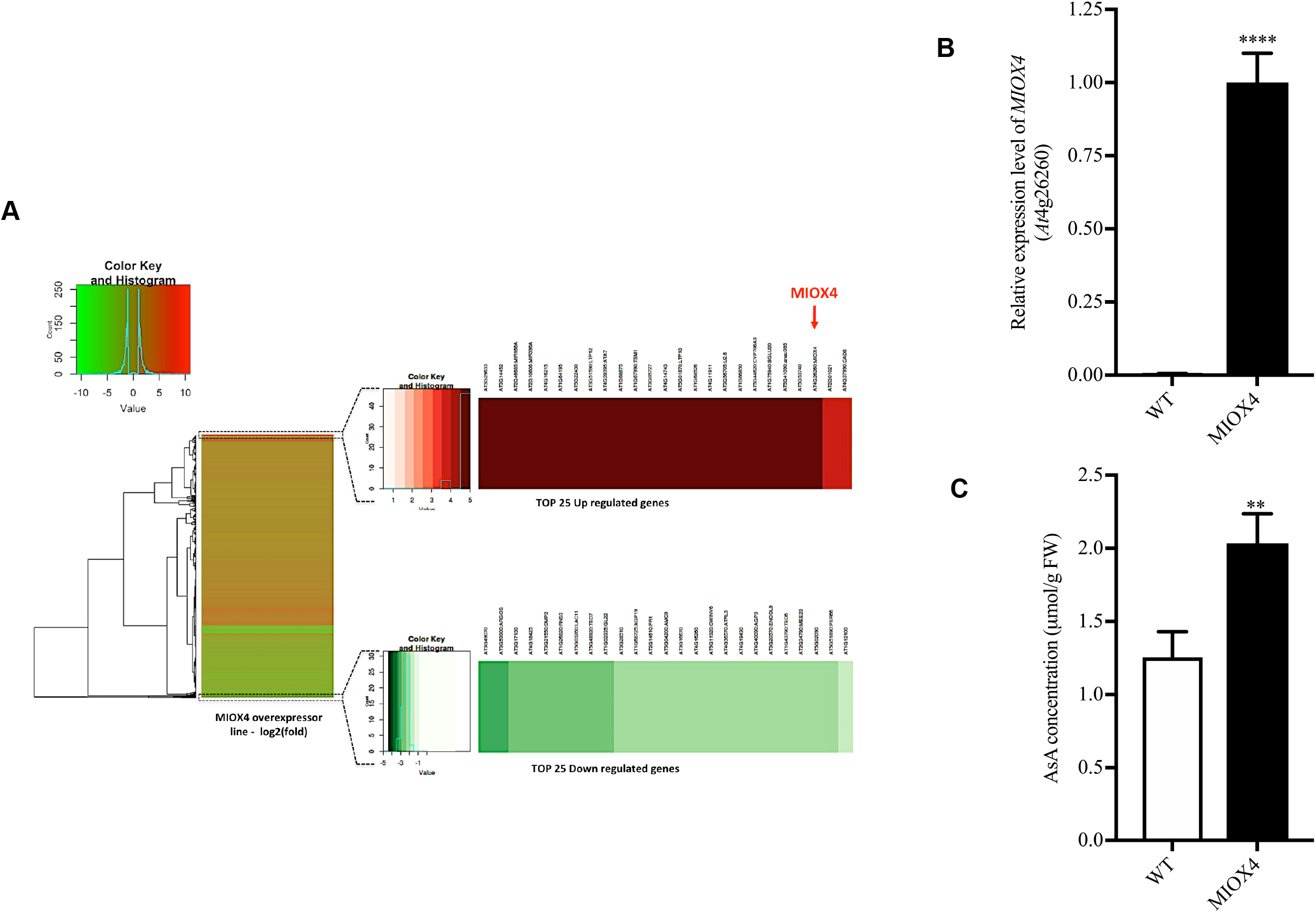
Global effects of the overexpression (OE) of *At*MIOX4 at the transcriptomic level. A) *MIOX4* was the highly up-regulated among differentially expressed transcripts shown in Heatmap and B) Validation of MIOX4 OE using RT-qPCR. Data was mean ± SEM (n=3) C) L-Ascorbate concentration was significantly higher in MIOX4 OE line compared to WT control. ** indicates p<0.01 **** indicated p-value<0.0001 at 0.05 significance level. Data was mean ± SEM (n=15).

### RNA-seq, assembly, and annotation

Foliar tissue from Arabidopsis *At*MIOX4 L21 plants and wild type controls was collected at developmental stage 5.0. High quality RNA was used to make six cDNA libraries, three from each genotype. These libraries were sequenced using the Illumina HiSeq 2500 Rapid Run flow cell with 1×50 bp paired end format. A total of 200,736,389 raw reads were obtained. The percentage of adapters in samples was up to 3.1%. The Trim-Galore script was used to remove the reads <40 bp in all samples. After removal of adapters and low quality reads, 185,235,265 bp of high quality reads were obtained. Reads that achieved a high Phred score (>38) indicating good quality sequences were used for the assembly. To assess the sequencing quality, the reads were mapped to the *A. thaliana* genome (TAIR10) using TopHat (Table 1). The functional gene ontology annotation of transcripts was performed using the Blast algorithm. The quality and mapping of the RNA-Seq data obtained from Illumina sequencing was good for transcriptome analysis of the *At*MIOX4 OE line.

**Table 1.** Quality analysis, trimming and aligning of the paired end reads using FastQC, Trimgalore and TopHAT.

| Sample | Total number of reads | Sequence length (bp) | Average Phred's value (~) | Number of sequences after filtering adapters (bp) | Nos. of reads after quality trimmed (bp) | Mapped left reads (bp) | Mapped right reads (bp) | Aligned pairs (bp) |
| --- | --- | --- | --- | --- | --- | --- | --- | --- |
| MIOX4 1_1 | 26210169 | 50 | 38 | 51,395,518 | 50,873,418<br>(99.9% of input) | 25,405,589<br>(97.9% of input) | 25,408,030<br>(97.9% of input) | 24,868,281<br>(95.8% of input) |
| MIOX4 1_2 | 25957145 | 50 | 38 |  |  |  |  |  |
| MIOX4 2_1 | 13523134 | 50 | 38 | 26,485,319 | 25,601,155<br>(99.8% of input) | 12,593,890<br>(94.2% of input) | 12,593,630<br>(94.2% of input) | 11,859,664<br>(88.6% of input) |
| MIOX4 2_2 | 13373369 | 50 | 38 |  |  |  |  |  |
| MIOX4 3_1 | 20571197 | 50 | 38 | 40,368,237 | 39,894,120<br>(99.9% of input) | 19,975,739<br>(98.0% of input) | 19,978,910<br>(98.0% of input) | 19,570,871<br>(95.9% of input) |
| MIOX4 3_2 | 20392195 | 50 | 38 |  |  |  |  |  |
| WT1_1 | 13878468 | 50 | 38 | 27,177,319 | 27,103,414<br>(99.9% of input) | 13,448,427<br>(97.9% of input) | 13,451,008<br>(97.9% of input) | 13,171,252<br>(95.9% of input) |
| WT 1_2 | 13734031 | 50 | 38 |  |  |  |  |  |
| WT 2_1 | 15061013 | 50 | 38 | 29,974,773 | 29,144,034<br>(99.9% of input) | 14,491,037<br>(97.2% of input) | 14,493,518<br>(97.2% of input) | 14,082,720<br>(94.42% of input) |
| WT 2_2 | 14913760 | 50 | 38 |  |  |  |  |  |
| WT 3_1 | 11630542 | 50 | 38 | 23,121,908 | 12,619,124<br>(99.9% of input) | 11,230,085<br>(97.7% of input) | 11,232,952<br>(97.8% of input) | 10,977,404<br>(95.5% of input) |
| WT 3_2 | 11491366 | 50 | 38 |  |  |  |  |  |

### MIOX4 is highly expressed in the AtMIOX4 line

Differentially expressed transcripts (DETs) in the *At*MIOX4 OE line were identified by comparison with the transcriptome of WT control. Genes with an adjusted p-value FDR <0.05 and a log_2_ fold change value of 2 were assigned as differentially expressed. As a result, 947 up-regulated transcripts and 357 down-regulated transcripts were differentially expressed in the *At*MIOX4 OE line compared to the control (Figure 3A). According to the functional annotation of DETs, most of the differential transcripts have been proven to be correlated to different biological processes related to developmental regulation and response to stresses in plants. The majority of up-regulated transcripts fall under the response to stimulus biological process. These include response to external stimulus, response to organic substance, response to endogenous stimulus, response to auxin stimulus, response to abscisic acid stimulus, response to jasmonic acid stimulus, response to stress, response to water deprivation, response to cold stress, and response to salt stress (Supplementary Table 4). On the other hand, the majority of down-regulated transcripts were grouped under biological processes as response to chemical stimulus, response to stimulus and response to stresses (Supplementary Table 5). We identified transcripts belonging to the precursors of three microRNAs (*At*3g63375, *At*2g46685, *At*2g10606), a transcript related with *TSM1* (*At*1g67990), *TRNG.1* (*At*Cg00100), and snoRNA (*At*1g32385, *At*1g20015) involved in cell regulation that were exclusively expressed in the *At*MIOX4 OE line. Similarly, transcripts related to stress responses such as lipid transfer proteins related to pathogenesis (*At*4g22513, *At*3g51590, *At*4g28395, *At*1g66850, *At*5g01870), an extensin family protein (*At*5g22430), a protease inhibitor (*At*4g22513), a guard cell specific transmembrane protein (*At*3g14452), a defensing-like family member protein (*At*1g64195, *At*3g05727), a stay-green like protein (*At*4g11911), a cytochrome P450 family protein (*At*5g44620), a β-glucosidase 20 (*At*1g75940), a NAC domain containing protein (*At*5g41090), and hypothetical proteins (*At*3g29633, *At*5g53740, *At*1g68526, *At*1g68875, *At*4g16240, *At*4g16215, *At*3g15534) were among the top 25 up-regulated genes in the *At*MIOX OE line (Table 2). On the other hand, development related genes such as a plant self-compatibility protein S1 family (*At*3g24065), an aspartyl protease family protein (*At*5g48430), a putative non-LTR retroelement reverse transcriptase (*At*2g07767; *At*3g49330), a plant invertase/pectin methylestarase inhibitor family protein (*At*5g12880; *At*3g17130), transmembrane proteins (*At*3g49070; *At*4g18425; *At*3g21550), *GASA10* (*At*5g59845), *ARGOs* (*At*3g59900), *IDL3* (*At*5g09805), *RNS3* (*At*1g26820), *LAC11* (*At*5g03260), *AGP19* (*At*1g68725) and stress response genes such as *PR1* (*At*2g14610), *GATA19* (*At*4g36620), and *MRD1* (*At*1g53480) were down-regulated in the *At*MIOX4 OE line compared to the control (Table 3). Additionally, amino acid metabolism genes lysine-ketoglutarate reductase/saccharopine dehydrogenase functional protein (*At*4g33150), involved in lysine degradation and induced by jasmonate (JA), and arginase (*At*4g08870), involved in defense response and induced by methyl jasmonate (MeJA), were abundant in *At*MIOX4 OE line compared to WT control (Supplementary Table 1-3; Figure 3A). The *MIOX4 (At*4g26260) gene was significantly over-expressed in the *At*MIOX4 transcriptome compared to the WT control, as shown by RT-qPCR (Figure 3B).

**Table 2.** List of top 25 up-regulated genes in *At*MIOX4 OE line compared to WT control

| Gene id | Gene | Log2<br>(fold_change) | TAIR information |
| --- | --- | --- | --- |
| *At3g29633* | - | inf | Encodes hypothetical protein |
| *Atcg00100* | *TRNG.1* | inf | Encodes tRNA-gly |
| *At1g32385* | - | inf | snoRNA-rRNA modification |
| *At3g14452* | - | inf | Encodes hypothetical transmembrane protein |
| *At1g20015* | - | inf | snoRNA-rRNA modification |
| *At3g63375* | *MIR167B* | inf | Targets ARF6 and ARF8 (auxin response factors-ARF) |
| *At2g46685* | *MIR166A* | inf | Targets several HD-ZIPIII family members |
| *At2g10606* | *MIR396A* | inf | Targets several GRF family members. |
| *At4g16240* | - | inf | Encodes hypothetical protein |
| *At4g16215* | - | inf | Encodes hypothetical protein |
| *At1g64195* | - | inf | Encodes a defensin-like (DEFL) family protein.. |
| *At3g15534* | - | inf | Encodes hypothetical protein |
| *At5g22430* | - | inf | Encodes pollen Ole e1 allergen and extensin family protein. |
| *At4g22513* | - | inf | Encodes a protease inhibitor/seed storage/LTP family protein |
| *At3g51590* | *LTP12* | inf | Encodes lipid transfer protein 12 |
| *At4g28395* | *ATA7* | inf | Encodes lipid transfer protein |
| *At1g68875* | - | inf | Encodes hypothetical protein |
| *At1g67990* | *TSM1* | inf | Tapetum specific methyl transferase |
| *At3g05727* | - | inf | Encodes a defensin-like (DEFL) family protein |
| *At4g14743* | - | inf | Encodes reverse transcriptase-related family protein |
| *At5g01870* | *LTP10* | inf | Predicted to encode a pathogenesis-related protein |
| *At1g68526* | - | inf | Encodes hypothetical protein |
| *At4g11911* | - | inf | Encodes stay-green-like protein |
| *At3g56705* | *U2.6* | inf | Encodes U2 small nucleolar RNA6 |
| *At1g66850* | - | inf | Encodes bifunctional inhibitor/lipid-transfer protein |

**Table 3.**
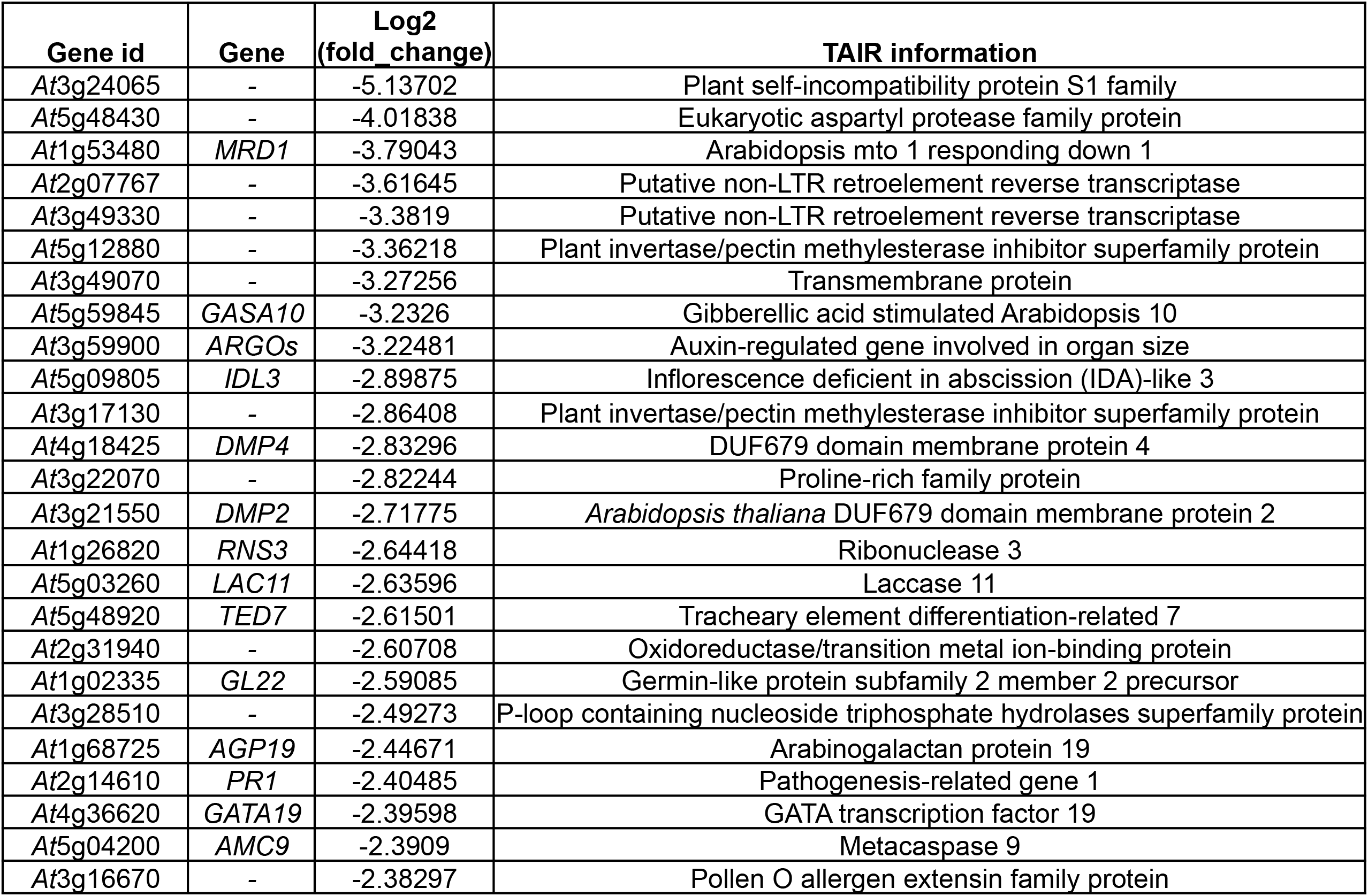
List of top 25 down-regulated genes in *At*MIOX4 OE line compared to WT control

| Gene id | Gene | Log2<br>(fold_change) | TAIR information |
| --- | --- | --- | --- |
| At3g24065 | - | -5.13702 | Plant self-incompatibility protein S1 family |
| At5g48430 | - | -4.01838 | Eukaryotic aspartyl protease family protein |
| At1g53480 | *MRD1* | -3.79043 | Arabidopsis mto 1 responding down 1 |
| At2g07767 | - | -3.61645 | Putative non-LTR retroelement reverse transcriptase |
| At3g49330 | - | -3.3819 | Putative non-LTR retroelement reverse transcriptase |
| At5g12880 | - | -3.36218 | Plant invertase/pectin methylesterase inhibitor superfamily protein |
| At3g49070 | - | -3.27256 | Transmembrane protein |
| At5g59845 | *GASA10* | -3.2326 | Gibberellic acid stimulated Arabidopsis 10 |
| At3g59900 | *ARGOs* | -3.22481 | Auxin-regulated gene involved in organ size |
| At5g09805 | *IDL3* | -2.89875 | Inflorescence deficient in abscission (IDA)-like 3 |
| At3g17130 | - | -2.86408 | Plant invertase/pectin methylesterase inhibitor superfamily protein |
| At4g18425 | *DMP4* | -2.83296 | DUF679 domain membrane protein 4 |
| At3g22070 | - | -2.82244 | Proline-rich family protein |
| At3g21550 | *DMP2* | -2.71775 | *Arabidopsis thaliana* DUF679 domain membrane protein 2 |
| At1g26820 | *RNS3* | -2.64418 | Ribonuclease 3 |
| At5g03260 | *LAC11* | -2.63596 | Laccase 11 |
| At5g48920 | *TED7* | -2.61501 | Tracheary element differentiation-related 7 |
| At2g31940 | - | -2.60708 | Oxidoreductase/transition metal ion-binding protein |
| At1g02335 | *GL22* | -2.59085 | Germin-like protein subfamily 2 member 2 precursor |
| At3g28510 | - | -2.49273 | P-loop containing nucleoside triphosphate hydrolases superfamily protein |
| At1g68725 | *AGP19* | -2.44671 | Arabinogalactan protein 19 |
| At2g14610 | *PR1* | -2.40485 | Pathogenesis-related gene 1 |
| At4g36620 | *GATA19* | -2.39598 | GATA transcription factor 19 |
| At5g04200 | *AMC9* | -2.3909 | Metacaspase 9 |
| At3g16670 | - | -2.38297 | Pollen O allergen extensin family protein |

We measured elevated foliar AsA content in the *At*MIOX4 OE (1.63 fold) compared to the control (Figure 3C), confirming our previous reports that over-expression of *At*MIOX4 leads to higher AsA content in Arabidopsis both in the absence and in response to abiotic stresses (Lorence et al., 2004; Lisko et al., 2013; Yactayo-Chang, 2011). The transcripts only expressed in the *At*MIOX4 OE line might have multiple roles in plant physiology, such as development, the oxidation-reduction process, carbohydrate and glucosinolate metabolism, and defense response against fungus, which could be vital for development and the stress tolerance/resistance phenotype observed in the *At*MIOX4 OE line. A selected number of the DET from each group were validated through RT-qPCR and those results are presented in the following sections.

### Primary metabolism is up-regulated in MIOX4 OE

The DETs were classified into different categories using the MapMan software (Usadel et al., 2009). Based on our results, the DETs were associated to primary metabolism, photosynthesis, biotic and abiotic stress, oxidative stress and secondary metabolism. Among the significantly over-expressed genes in the *At*MIOX4 OE line involved in primary metabolism, we identified transcripts related with sugar metabolism such as hexokinase like protein 1 (*HKL1*: *At*1g50460), α-amylase-like (*AMY1*: *At*4g25000), and cellulose synthase (*At*4g24000). In addition, transcripts related to sugar and starch metabolism were significantly up-regulated in *At*MIOX4 OE line compared to the control (Figure 4A). Interestingly, a starch degrading enzyme, α-amylase (*At*4g25000), was significantly up-regulated in the *At*MIOX4 transcriptome which was validated using RT-qPCR (Figure 4B). α-Amylase catabolizes starch and releases free glucose and dextrin. Therefore we proceeded to measure the intracellular pool of free glucose in the high AsA line using two different methods. As show in Figure 4C we found that the intracellular concentration of reducing sugars was significantly higher in the *At*MIOX4 OE line (41.64 ± 7.61 μM/g FW) compared to controls (23.13 ± 2.02 μM/g FW). In support of this, the intracellular concentration of glucose in the *At*MIOX4 OE line (31.92 ± 0.95 μM/g FW) was significantly higher compared to the one found in the control (19.61 ± 0.99 μM/g FW), as shown by a glucose specific enzymatic assay (Figure 4D). In summary, intracellular glucose concentration was 1.62 fold higher in *At*MIOX4 OE line compared to WT control. This indicates that the *At*MIOX4 OE line has elevated expression of α-amylase which leads to higher available intracellular glucose in these plants.

**Figure 4.**
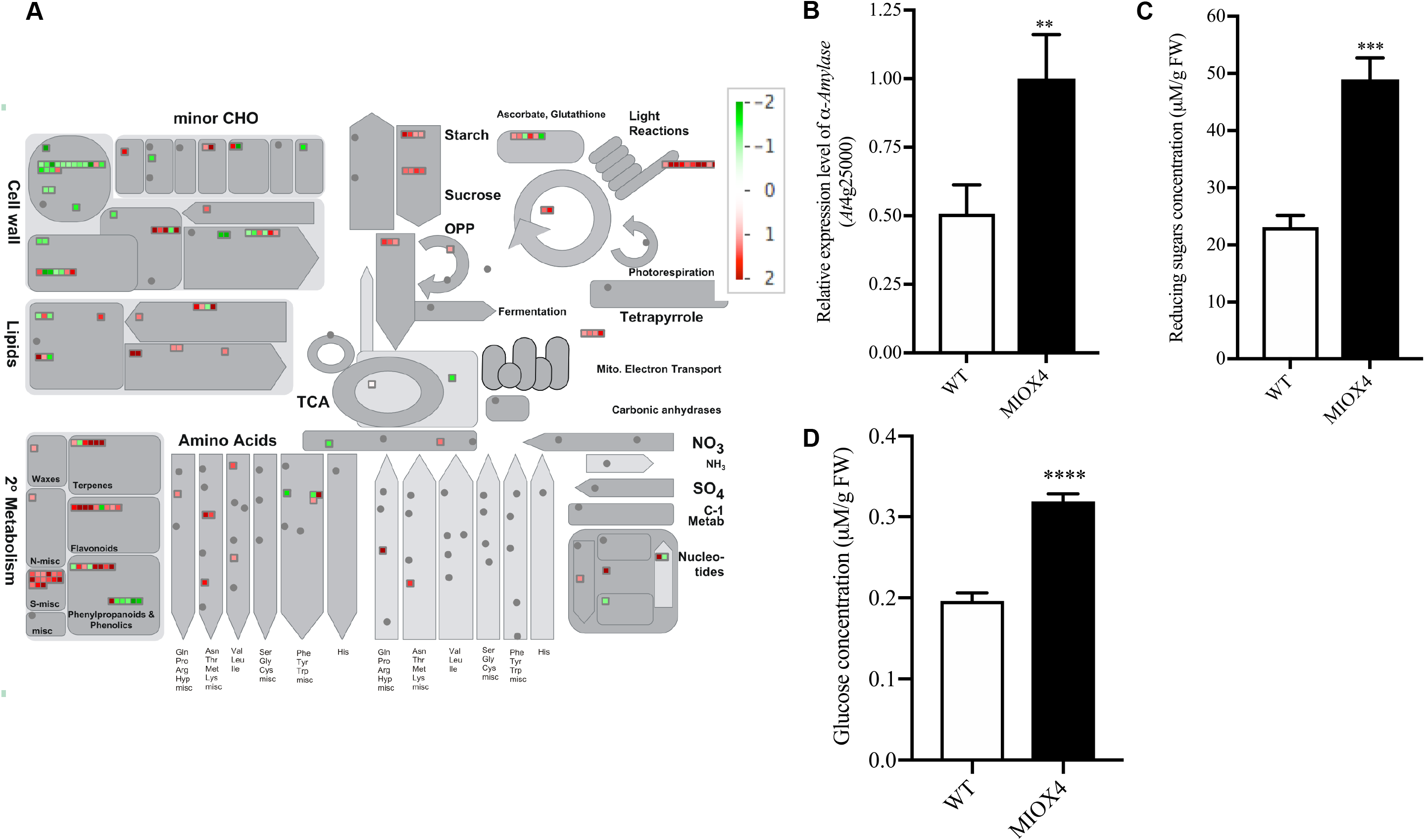
Intracellular glucose concentration was higher in MIOX4 OE line relative to wild type (WT) plants. A) The overall effect on physiological mechanism in MIOX4 OE compared to WT plant. Figure was obtained using MapMan 3.5.1 R2. (Up-regulated transcripts was indicated by red square and down-regulated transcripts were indicated by green square) B) α-amylase-like gene was significantly up-regulated in MIOX4 OE compared to WT plant shown by RT-qPCR. Data was mean ± SEM (n=3) C) Reducing sugars concentration were significantly higher in MIOX4 OE line compared to WT line shown by DNS assay. D) Glucose concentration was significantly higher in MIOX4 OE line compared to WT line shown by GOD-POD assay. ** indicated p<0.01, *** indicated p<0.001 and **** indicated p<0.0001 at 0.05 significance level. DNS=*Di*-nitosalicylic acid, GOD-POD=Glucose oxidase-Glucose peroxidase. Figures are Mean ± SEM of three replicates for RT-qPCR and six replicates for DNS and GOD-POD assay. Data was mean ± SEM (n=6)

### MIOX4 over-expression positively influences photosynthetic efficiency

Transcripts involved in light reactions of photosynthesis, as photosystem II related transcripts (*PSBA*-ATCG00020, *PSBC*-ATCG00280, *PSBB*-ATCG00680) and photosystem I related transcripts (*PSAJ*-ATCG00630, *PSAC*-ATCG01060, *NDHG*-ATCG01080) were significantly up-regulated in the *At*MIOX4 OE line (Figure 5A). In support of this, over-expression of photosystem II center related transcript (*PSBB*-*At*Cg00680) was validated with RT-qPCR (Figure 5B). To determine if the increased gene expression we observed is physiologically significant, we measured photosynthetic efficiency using a handheld device (Kuhlgert et al., 2016). As illustrated in Figures 5C-D photosystem II efficiency and proton motive force (vH^+^) were significantly higher in the *At*MIOX4 OE line compared to WT control at 21, 23, 26 and 28 days after germination. Taking together these results show that the *At*MIOX4 OE line is photosynthetically more efficient than its WT counterpart.

**Figure 5.**
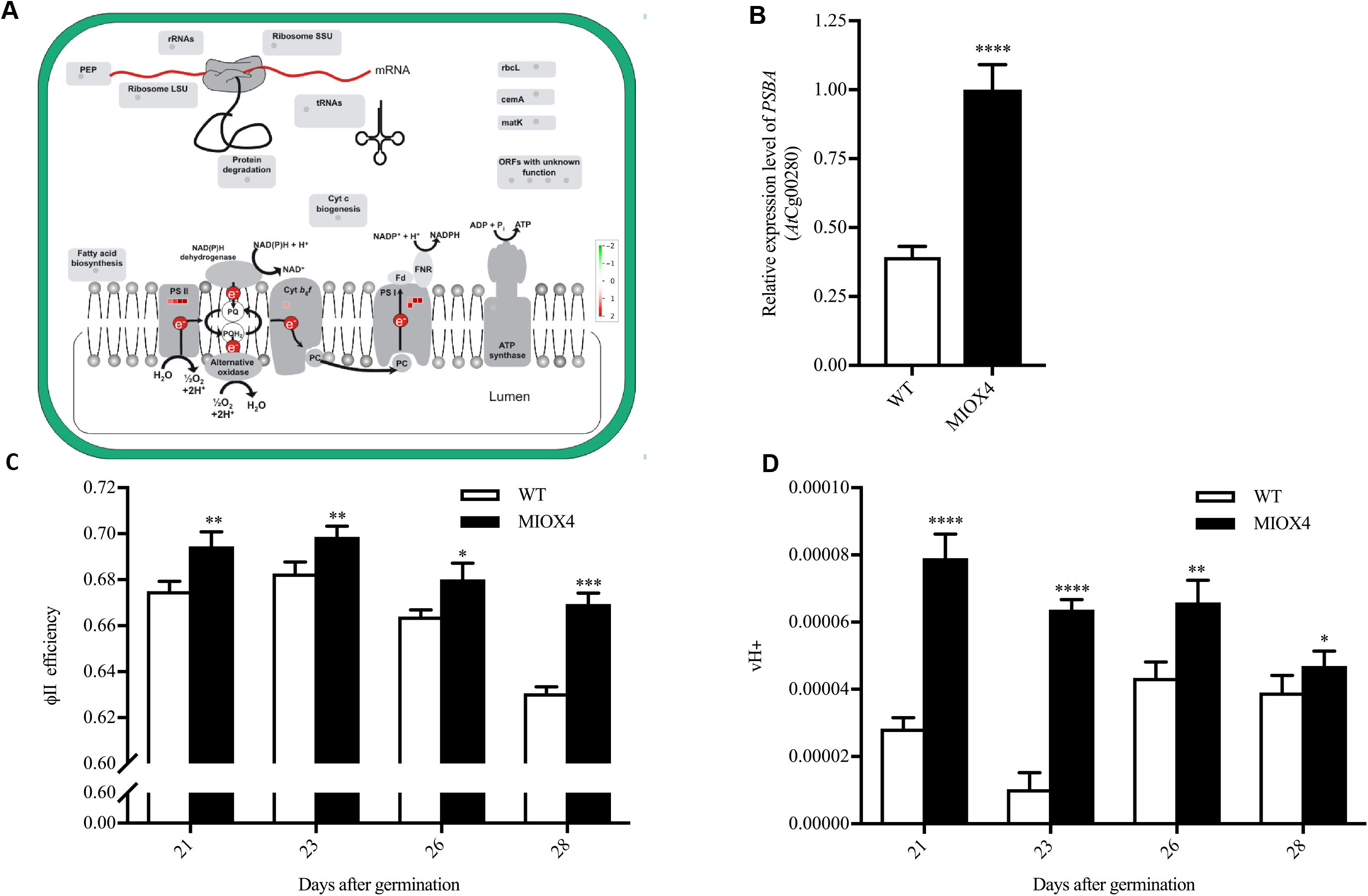
The high ascorbate MIOX4 OE line is photosynthetically more efficient. A) Transcripts related to photosystem I and II (PSI, PSII) were up-regulated (indicated by red square) in MIOX4 OE line compared to WT line. The figures were obtained using MapMan V3.5.1R2 B) Overexpression of transcripts related to PSII center was validated using RT-qPCR. Data was mean ± SEM (n=3) C),D) Photosystem II efficiency (ΦII efficiency) and protonmotive force (vH^+^) were significantly higher in MIOX4 OE line compared to WT line from 21 to 28 days after germination. Data was mean ± SEM (n=15). ** indicated p<0.01, *** indicated p<0.001 and **** indicated p<0.0001 at 0.05 significance level.

### MIOX4 over-expression leads to increase auxin

Next, we investigated the role of phytohormones in the enhanced growth and abiotic stress tolerance phenotype recorded for the *At*MIOX4 OE line. Analysis of the transcriptome data indicate that genes involved in auxin biosynthesis (*YUCCA6: At5*g25620, *NIT2: At*3g44300), auxin-amino acid conjugate hydrolase (*ILL6: At*1g44350), auxin transport (auxin efflux carrier family protein), and small auxin up RNAs (*SAUR2: At*4g34780, *SAUR4: At*4g34800, *SAUR36: At*2g45210) were up-regulated in *At*MIOX4 OE line compared to WT controls, whereas auxin response small RNA (*SAUR53: At*1g19840) and auxin regulated organ size transcript (*ARGOs: At*3g59900) were down-regulated in *At*MIOX4 OE line compared to WT (Figure 6A). The differential expression of auxin biosynthesis (*YUCCA6, NIT2*), auxin-amino acid conjugate hydrolase (*ILL6*), and transcripts involved in regulation of organ size (*ARGOs*) was confirmed via RT-qPCR (Figure 6B).

**Figure 6.**
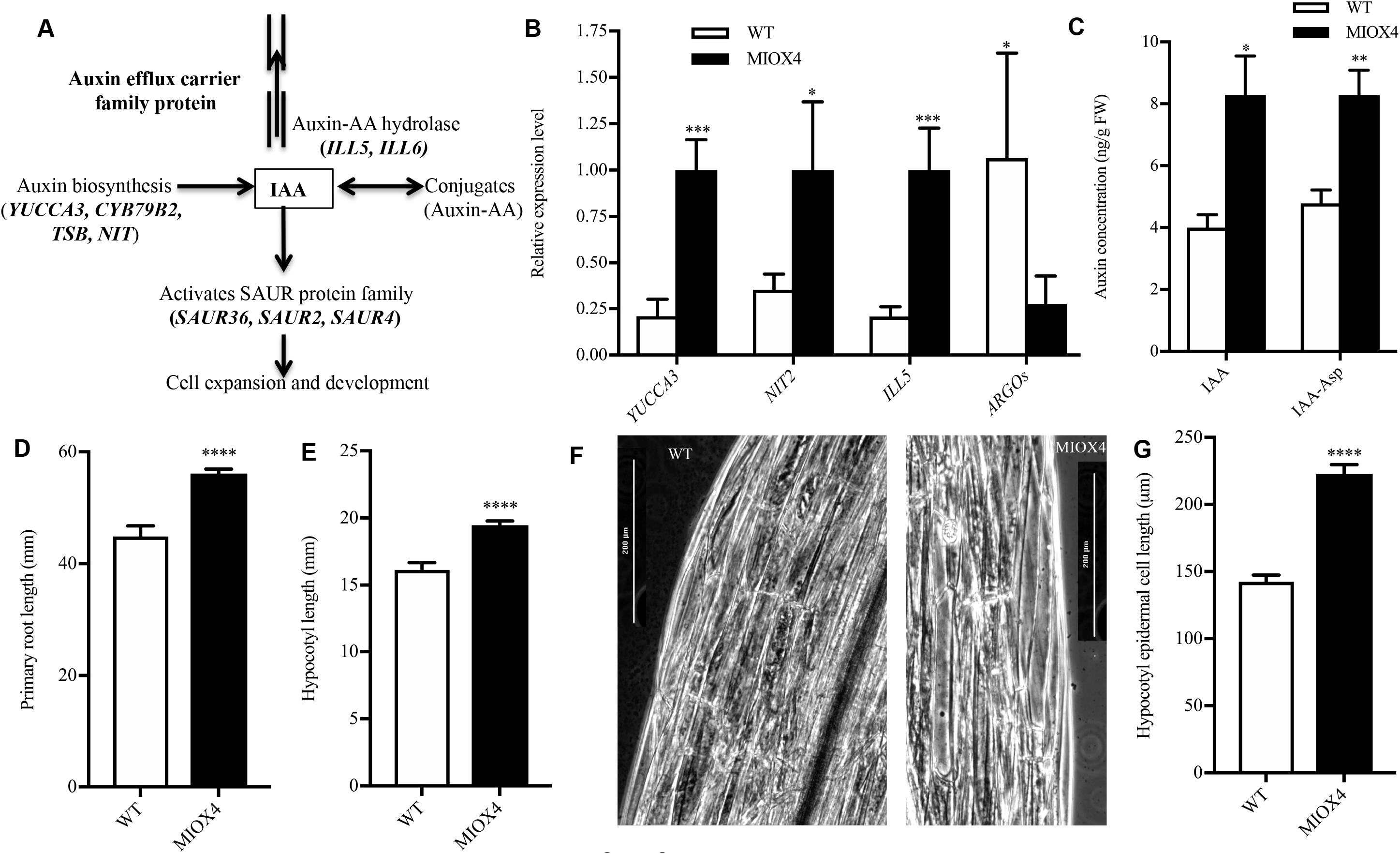
Increased level of auxin in high ascorbate MIOX4 OE line are biologically more relevant. A) Transcripts related to auxin biosynthesis, auxin-aminoacid conjugate hydrolases, auxin transport and auxin response small RNA were up-regulated in MIOX4 OE transcriptome compared to WT line. B) Transcripts related to auxin metabolism were validated using RT-qPCR. Data was mean ± SEM (n=3) C) Active auxin (IAA) was significantly higher but inactive auxin (IAA-Asp) was similar in MIOX4 OE line compared to WT line as shown by LC-MS/MS. Data was mean ± SEM (n=5) D), E) Hypocotyl length grown in dark conditions and primary root length grown in light condition were significantly higher in MIOX4 line compared to WT line in 9 days after germination. Data was mean ± SEM (n=20) F), G) Hypocotyl epidermal cell length were significantly higher in MIOX4 OE line compared to WT line. ** indicated p<0.01, *** indicated p<0.001 and **** indicated p<0.0001 at 0.05 significance level. Data was mean ± SEM (n=30)

Next we determined the intracellular level of auxin and its conjugated form using LC-MS/MS. The intracellular concentrations of the active auxin (IAA) and inactive auxin (IAA-Ile) were significantly higher in *At*MIOX4 OE line compared to WT (Figure 6C). To understand the effect of elevated auxin in *At*MIOX4 OE line, next we measured the elongation of the primary root in light conditions and the hypocotyl length/hypocotyl epidermal cell length in dark conditions in the *At*MIOX4 and controls. As shown in Figure D-E, the primary root of the high AsA line was longer than the control when plants grew under light conditions, and their hypocotyl was longer when plants grew in the dark. Next, we investigated whether the hypocotyl elongation in the *At*MIOX4 OE line was due to increase in cell number or cell length using phase contrast microscopy. We found that hypocotyl epidermal cell length was significantly elongated in the *At*MIOX4 OE line compared to WT controls (Figure 6F-G). Subsequently, we took advantage of the auxin sensor line R2D2 developed by Liao et al (2015) to monitor *in vivo* auxin levels in our high AsA line. For this, crosses between the R2D2 and the *At*MIOX4 line (both lines are present in the *A. thaliana* Col-0 ecotype) were made. Positive crosses were selected using kanamycin as a selectable marker and PCR to confirm the presence of the transgenes of interest. Two positive crosses and controls were used for the *in-vivo* auxin visualization and semi-quantification. The R2D2 line expresses lower ntdTomato fluorescence (red) in the presence of normal auxin level. When there is a higher level of auxin, there is a higher level of ntdTomato fluorescence, compared to n3 × DII-Venus (green) expression which was prominent in both shoots (Figure 7A-B) and roots (Figure 7D-E). As illustrated in Figure 7 the ratio of ntdTomato fluorescence to n3X DII Venus fluorescence was significantly higher in the crosses compared to R2D2 control in both shoots (Figure 7C) and roots (Figure 7F). We did not find evidence of differentially regulated transcripts involved in cytokinin metabolism in the *At*MIOX4 OE line. The level of active cytokinin (*cis*-zeatin), isomeric and relatively less active cytokinin (*trans*-zeatin), and cytokinin conjugate (zeatin-riboside) levels were similar in *At*MIOX4 OE line compared to WT control as data from LC-MS/MS shows (Figure 8A-C). Taken all together, these results indicate that there are elevated levels of auxin due to the transcriptional up-regulation of auxin metabolism in the *At*MIOX4 OE line and that the elevated auxin levels observed in the *At*MIOX4 OE line are functionally relevant. Our observations support a major role for auxin in cell elongation leading to the higher mass phenotype of the high AsA line.

**Figure 7.**
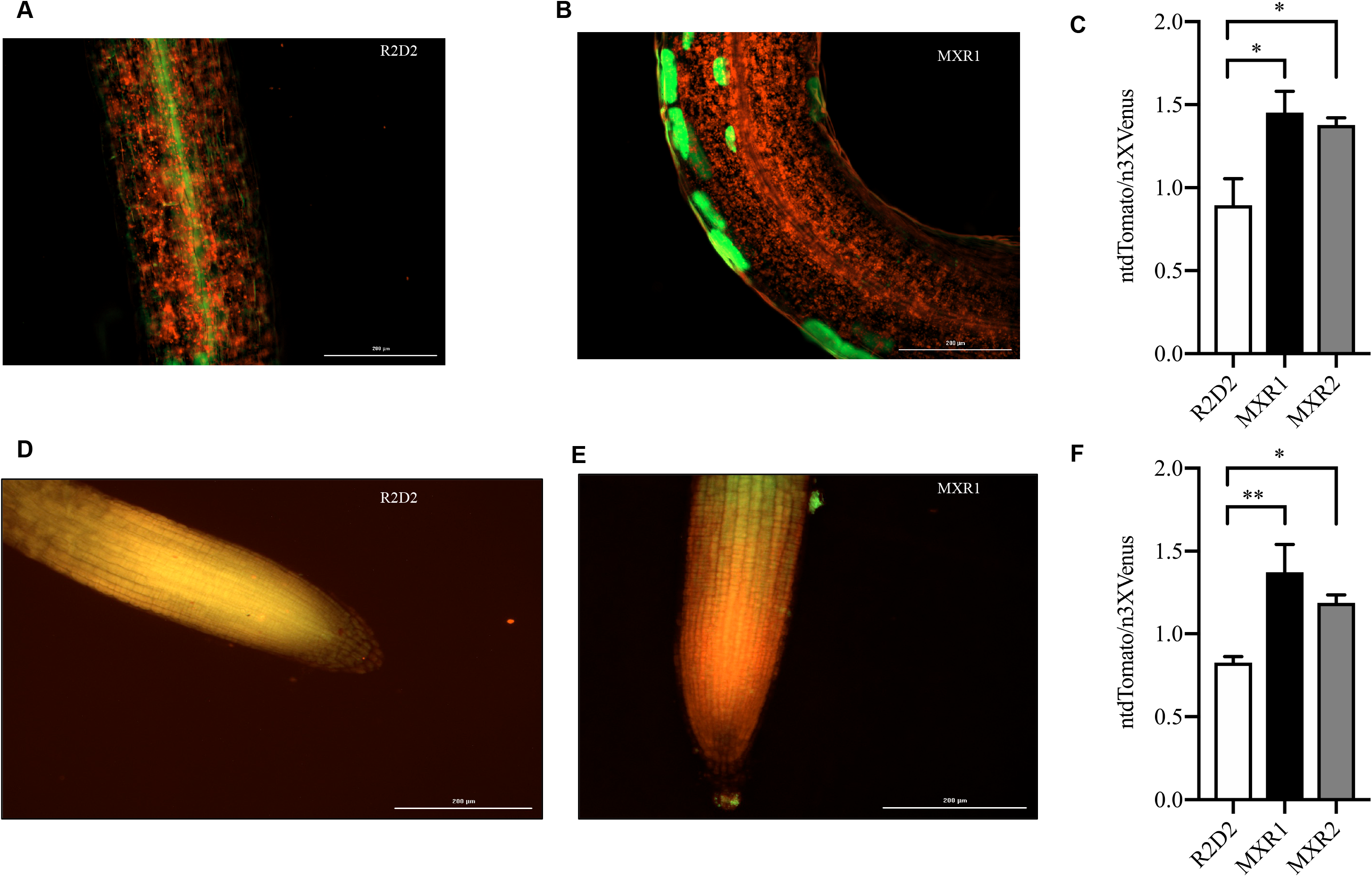
*In-vivo* visualization and semi-quantification of auxin response. A), B), C)The ntdTomato expression was significantly higher i MIOX4 and R2D2 crosses compared to R2D2 control in shoots. D), E), F) The ntdTomato expression was significantly higher in MIOX and R2D2 crosses compared to R2D2 control in roots. ntdTomato=red, DII-venus=green. Data was mean ± SEM (n=4). Scale bars=200μm). Data was analyzed using one-way ANOVA and multiple comparisons was performed comparing crosses with R2D2 lines using Dunnett’s multiple comparisons test at 0.05 significance level. * indicates p<0.05 and ** indicates p<0.01.

**Figure 8.**
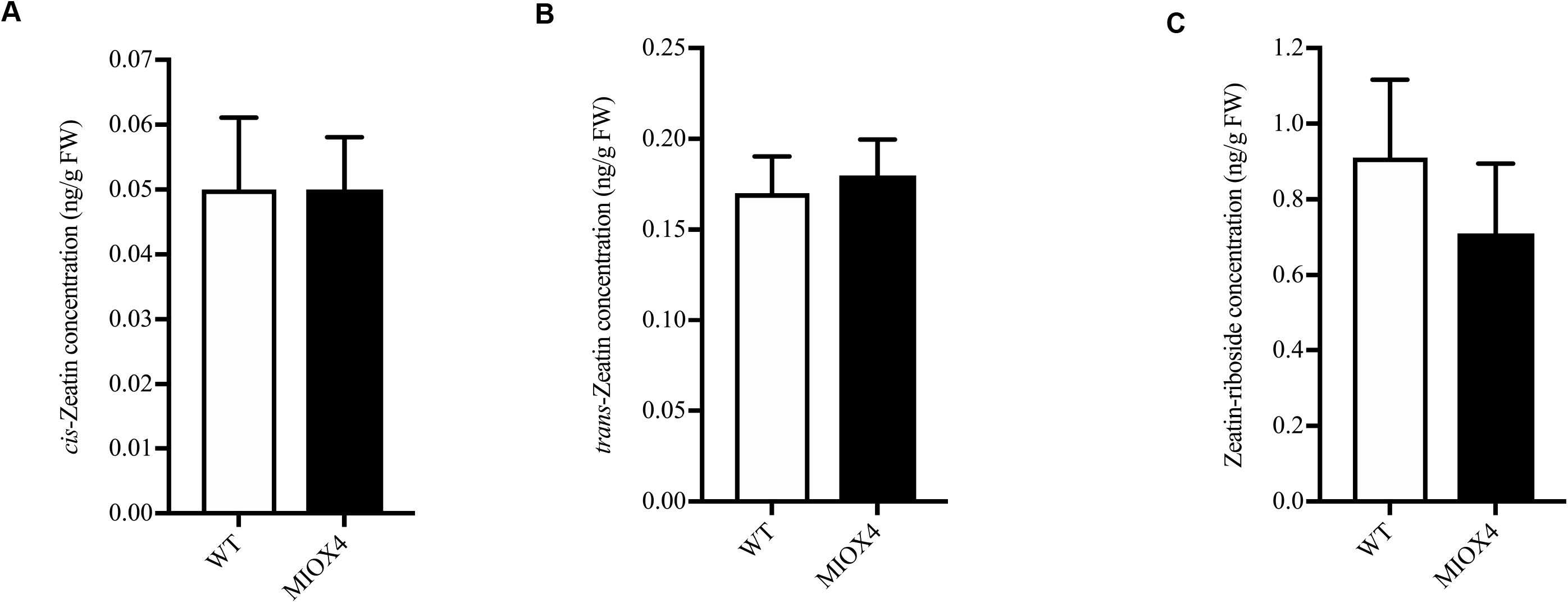
Cytokinin metabolism were unaffected by MIOX4 overexpression. A) *trans*-zeatin B) *cis*-zeatin and C) zeatin-riboside levels were similar in MIOX4 OE line compared to WT line as shown by LC-MS/MS at 0.05 significance level. Data was mean ± SEM (n=10)

### MIOX4 over-expressers tolerate abiotic stresses independently of abscisic acid up-regulation

We also examined some of the possible molecular mechanisms mediating the abiotic stress tolerance phenotype in *At*MIOX4 OE line. We identified abiotic stress response-related based on the expression profiles of the DETs transcripts using MapMan. Among the statistically significant transcripts we found that heat stress response transcripts 17.8KDa class I heat shock protein (*HSP17.8-CI*-*At*1g07400), 17.4 KDa class III heat shock protein (*HSP17.4-CIII*-*At*1g54050), cold regulated protein 15a (*COR15A*-*At*2g42540) and the drought induced protein (*ATDI21*-*At*4g15910) were significantly up-regulated in *At*MIOX4 OE line compared to WT control (Figure 8A).

We also looked at whether levels of abscisic acid were altered in the *At*MIOX4 OE line compared to controls using LC-MS/MS. Abscisic acid levels were similar in *At*MIOX4 OE and the WT line (Figure 8C). We then used Genevestigator to find signature genes that were over-expressed in publically available abiotic stresses experiments. Genevestigator analysis of the top 200 up-regulated (log_2_fold>1.75) and down-regulated transcripts (log_2_fold<−1.25) was compared against signature expression transcripts from 50 similar perturbations. We have found that cinnamyl alcohol dehydrogenase (*ATCAD8*: *At*4g37990), N-acetyl transferase activity 1 (*At*2g39030), lactolyl glutathione lyase (*GLYI7*:*At*1g80160), bHLH family protein (*At*1g10585) and NAC domain containing protein 19 (*NAC019*:*At*1g52890) were up-regulated in both, the set of abiotic stress perturbation experiments and in the *At*MIOX4 OE line (Supplementary Figure 4). Taken together, these results point out to an important role for a conserved module of stress response genes in the abiotic stress tolerance phenotype observed in *At*MIOX4 OE line.

### High AsA exerts a contrasting regulation on glutathione reductase and ascorbate peroxidase genes

Antioxidants maintain the homeostasis of ROS in plants. We looked for genes involved in the antioxidant cycle network in the high AsA line to identify possible crosstalk within the redox network. We found that superoxide dismutase expression was similar in *At*MIOX4 OE line compared to WT control (Figure 9A). Glutathione reductase 480 (*GRX480*: *At*1g28480), which regulates the protein redox state, and dehydroascorbate reductase 1 (*DHAR1*: *At*1g19570), involved in recycling of reduced ascorbate, were up-regulated while peroxisomal ascorbate peroxidase 5 (p*APX5*: *At*4g35970), involved in scavenging ROS, was down-regulated in the *At*MIOX4 OE line compared to WT control. The expression of *APX5* and *GRX480* was validated using RT-qPCR (Figure 9B). Interestingly, expression of genes involved in multiple pathways of L-ascorbate biosynthesis were similar in the *At*MIOX4 OE and the control, while expression of *DHAR1*, involved in recycling of reduced ascorbate, was up-regulated (Supplementary Figure 1). Down-regulation of peroxisomal *APX5* leads to significantly higher reactive oxygen species (H_2_O_2_) in *At*MIOX4 OE compared to its WT counterpart (Figure 9C). Taking together these results indicate that high *At*MIOX4 OE leads to higher ROS concentration due to down-regulation of peroxisomal *APX* gene. Additionally, high AsA leads to up-regulation of glutathione peroxidase and *DHAR1* genes which maintain the ROS homeostasis in *At*MIOX4 OE line.

**Figure 9.**
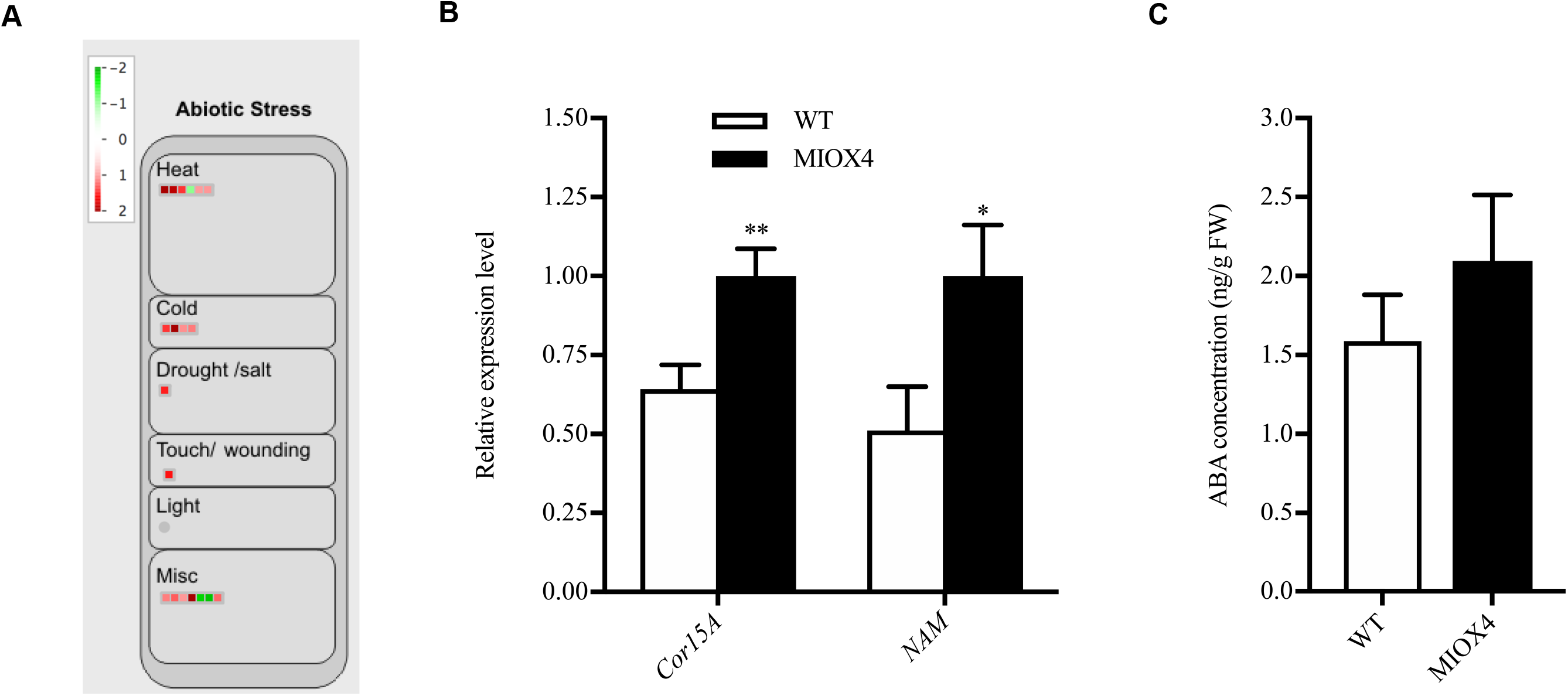
Abiotic stress response genes were differentially expressed in high ascorbate line. A) The heat, cold and drought stress response transcripts were up-regulated in MIOX4 OE line compared to WT line. B) Validation of over-expression of cold regulated protein (*CoR15a*) and *NAM* transcription factor using RT-qPCR. Data was mean ± SEM (n=3) C) The abscisic acid level remained similar in MIOX4 OE line and WT line as shown by LC-MS/MS. * indicated p<0.05 at 0.05 significance level. Data was mean ± SEM (n=10)

**Figure 10.**
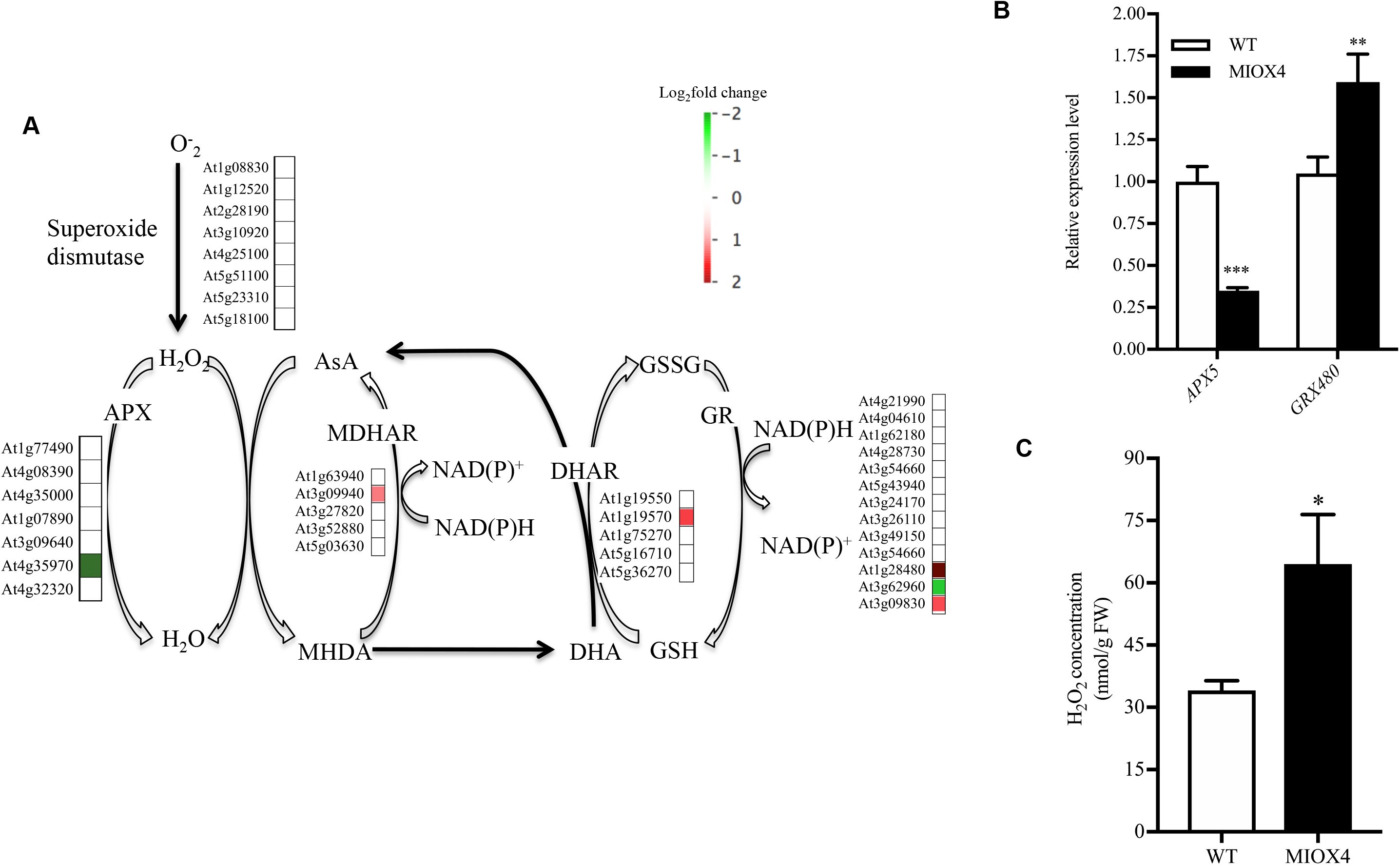
Antioxidants crosstalk between ascorbate, glutathione, superoxide dismutase, peroxidase and catalase. A) Ascorbate recycling enzyme *DHAR1*, Glutathione reductase(*GRX480*) were significantly up-regulated whereas ascorbate peroxidase 5 (*APX5*) were significantly down-regulated in MIOX4 OE transcriptome compared to WT line. B) RT-qPCR validation of up-regulated and down-regulated transcripts involved in ascorbate-antioxidant crosstalk. Data was mean ± SEM (n=3). C) H2O2 accumulation is higher in foliar tissue of MIOX4 OE line compared to WT line. Data was mean ± SEM (n=4). * indicated p<0.05, ** indicated p<0.01 and *** indicated p<0.001 at 0.05 significance level. APX=Ascorbate peroxidase, AsA=Ascorbate, GR=Glutathione reductase, GSH=Reduced glutathione, GSSG=Oxidized glutathione, H2O2=Hydrogen peroxide,

### High ascorbate plants seem ready to defend themselves against herbivores

Genes related to cell-wall metabolism, specialized metabolism, and defense hormone biosynthesis are important for the biotic stress response of plants. We found that genes involved in arabinogalactan metabolism and xyloglucan metabolism were down-regulated, whereas cellulose synthesis was up-regulated in *At*MIOX4 OE line compared to controls. Three members of the Arabinogalactan gene family, *AGP19*: *At*1g68725, *AGP3* (*At*4g40090) and *AGP14* (*At*5g56540) were down-regulated. Similarly, xyloglucan metabolism genes *XTH17* (*At*1g65310), *XTH25* (*At*5g57550) and *XTH10* (*At*2g14620) were down-regulated, whereas genes related to cellulose synthesis, *CSLA 10* (*At*1g24070), *CSLA 1* (*At*4g16590) and *CSLG2* (*At*4g24000) were up-regulated in *At*MIOX4 OE line compared to WT control.

Specialized metabolites are also an important line of defense in plants to combat stresses. Genes related to phenolic biosynthesis, O-methyl transferase family 2 protein (*At*1g76790), were up-regulated, while laccase biosynthesis genes, *LAC11* (*At*5g03260) and *LAC17* (*At*5g60020) were down-regulated in the *At*MIOX4 OE line compared to WT. The phenylpropanoid biosynthesis gene, *TSM1* (*At*1g67990) was only expressed in the *At*MIOX4 OE line, while *A. thaliana CAD-B2* (*At*4g37990) was up-regulated in *At*MIOX4 OE line compared to controls. Flavonoid metabolism related transcription *MYB 75* (*At*1g56650), which is involved in radical scavenging activity and anthocyanin metabolism, 2OG-Fe(II) oxygenase family proteins (*At*2g38240, *At*3g55970, *At*5g05600), and *AACT1* (*At*5g61168) were up-regulated in *At*MIOX4 OE line compared to controls. Similarly, terpene biosynthesis genes, *GGPS6* (*At*1g49530), *TPS10* (*At*2g24210), *LAS1* (*At*3g45310), and *ATTPS3* (*At*4g16740) were up-regulated in *At*MIOX4 OE line compared to WT.

Defense hormones, such as jasmonic acid (JA), salicylic acid (SA), and ethylene are critical to induce the systemic acquired defense response in plants. Genes related to JA biosynthesis and metabolism, *OPR3* (*At*2g06050) and *JR1 (At*3g16470) were up-regulated in the *At*MIOX4 OE line compared to WT control. Similarly, genes related to SA metabolism, *MES9* (*At*4g37150), involved in conversion of methyl salicylate to active SA, UDP*-* glucosyl transferase family protein (*At*1g05680), and *BSMT1* (*At*3g11480) were also up-regulated in the *At*MIOX4 OE line. JA response transcripts, *VSP2* (*At*4g11740), *VSP1* (*At*5g24780), and *PDF1.4* (*At*1g19610) were significantly up-regulated in the *At*MIOX4 OE line compared to WT. Similarly, we noted that ethylene response factors as *RAP2.6* (*At*1g43160), *RAP 2.6L* (At5g1330), and *RAP2.10* (*At*4g36900) were significantly up-regulated in the *At*MIOX4 OE line compared to WT (Figure 11A; Table 4). The expression of the signature genes that play a significant role in biotic stress response (*MYB27: At*3g53200, *OPR3, VSP2: At*4g11740 and *RAP2.6: At*1g43160) were validated using RT-qPCR (Figure 11B). *OPR3* regulates the biosynthesis of JA in plants. The over-expression of the *OPR3* leads to higher levels of JA and JA-Ile, while the metabolite intermediate 12-oxophytodieonate (OPDA) was similar in *At*MIOX4 OE line compared to WT control (Figure 11C-E). The over-expression of the genes related to specialized metabolites, such as phenylpropanoid, flavonoid, and terpene biosynthesis; up-regulation of genes involved in JA and SA biosynthesis; elevated level of JA and JA-Ile hormones; and expression of JA response genes, such as *VSP1, VSP2*, and *PDF 1.4* suggests that *At*MIOX4 OE plants are ready to defend themselves from herbivores attack.

**Figure 11.**
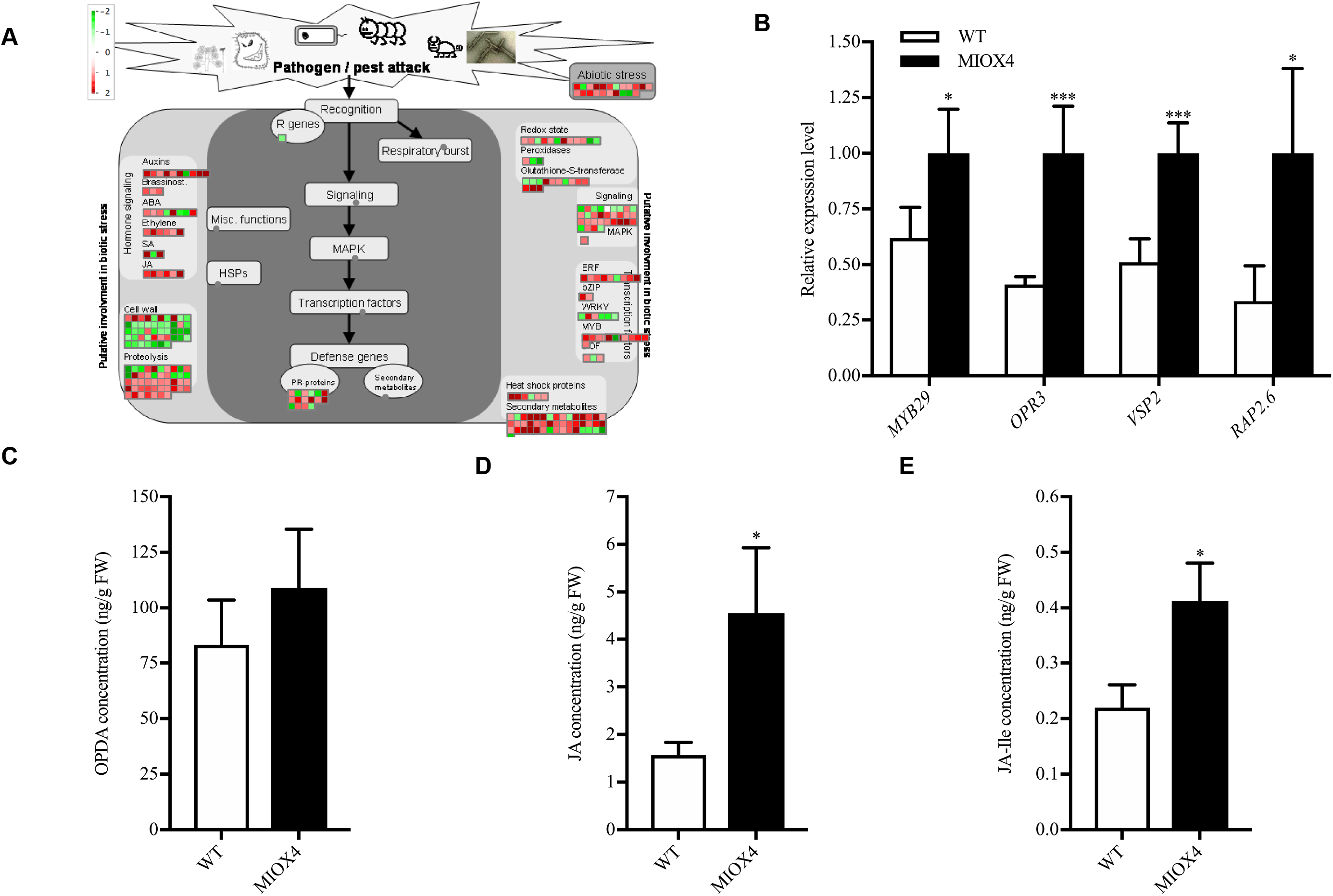
Biotic stress response transcripts are differentially expressed in high ascorbate MIOX4 OE line. A) Transcripts related to defense hormone biosynthesis, biotic stress response transcription factors, secondary metabolite biosynthesis were up-regulated and the transcripts related to cell wall metabolism were down-regulated in MIOX4 OE line compared to WT control. B) The up-regulation of transcripts involved in biotic stress response were validated using RT-qPCR. Data was mean ± SEM (n=3) C) Defense hormone Jasmonates (JA) and d) Jasmonic acid-Ile (JA-Ile) were significantly higher in MIOX4 OE line compared to WT control. Data was mean ± SEM (n=5). * indicated p<0.05, ** indicated p<0.01 and *** indicated p<0.001 at 0.05 significance level.

**Table 4.** Defense response genes up-regulated in *At*MIOX4 OE line relative to WT plants.

| Gene Id | Gene | Log2<br>(fold change) | TAIR information |
| --- | --- | --- | --- |
| *At1g67990* | *TSM1* | Inf | Encodes tapetum specific O-methyltransferase |
| *At4g37990* | *CAD-B2* | 3.85 | Encodes an aromatic alcohol:NADP+ oxidoreductase |
| *At1g56650* | *MYB 75* | 1.65 | Encodes MYB domain containing transcription factor |
| *At2g38240* | *JAO4* | 2.26 | Encodes 2-oxoglutarate/Fe(II)-dependent oxygenases |
| *At3g55970* | *JAO3* | 3.04 | Encodes 2-oxoglutarate/Fe(II)-dependent oxygenasees |
| *At5g05600* | *JAO2* | 2.38 | Encodes 2-oxoglutarate/Fe(II)-dependent oxygenases |
| *At5g61168* | *AACT1* | -1.75 | Encodes anthocyanin 5-aromatic acyltransferase 1 |
| *At1g49530* | *GGPS6* | 1.57 | Encodes mitochondrial-targeted geranyl geranyl pyrophosphate synthase 6 |
| *At2g24210* | *TPS10* | 3.39 | Encodes terpene synthase 10 |
| *At3g45310* | *LAS1* | 2.01 | Encodes cysteine protease superfamily protein |
| *At4g16740* | *ATPSO3* | 2.79 | Encodes an (E,E) -alpha-farnese synthase |
| *At2g06050* | *OPR3* | 1.76 | Encodes 12-oxophytodienate reductase |
| *At3g16470* | *JR1* | 2.04 | Encodes jasmonate-responsive 1 |
| *At4g37150* | *MES9* | 2.04 | Encodes methylesterase 9 |
| *At1g05680* | *UGT74E2* | 3.70 | Encodes uridine diphosphate glycosyltransferase 74E2 |
| *At3g11480* | *BSMT1* | 2.33 | Encodes SABATH methyltransferase |
| *At4g11740* | *VSP2* | 3.12 | Encodes vegetative storage protein 2 |
| *At5g24780* | *VSP1* | 3.15 | Encodes vegetative storage protein 1 |
| *At1g19610* | *PDF1.4* | 2.03 | Predicted to encode plant defensin family protein |
| *At1g43160* | *RAP2.6* | 1.77 | Encodes ethylene response factor subfamily B-4 of ERF/AP2 transcription factor family |
| *At5g13330* | *RAP 2.6L* | 2.28 | Encodes ethylene response factor subfamily B-4 of ERF/AP2 transcription factor family |
| *At4g36900* | *RAP2.10* | 1.46 | Encodes member of the DREB subfamily A-5 of ERF/AP2 transcription factor family |

### Genes from the glucosinolate biosynthetic pathway are upregulated in MIOX4 over-expressers

Glucosinolates are specialized metabolites present mostly in plants of the Brassicaceae family. It is well known that glucosinolates and their hydrolysis products act as defense compounds against herbivores and pathogens. Glucosinolates are derived from amino acids and are grouped into aliphatic, aromatic, and indolic glucosinolates based on their different side chain structures. We have found that genes involved in the glucosinolate -myrosinase system, flavin monoxygenase glucosinolate S-oxygenase 2 (*FMOGS-OX2*: *At*1g62540), alkenyl hydroxalkyl producing 2 (*AOP2*: *At*4g03060), MYB domain protein 76 (*MYB76*: *At*5g07700), myrosinase binding protein 1 (*MBP1*: *At*1g52040), cytochrome P450 *CYP83B1* (SUR2: *At*4g31500), and epithiospecifier protein (*ESP*: *At*1g54040) were elevated in the *At*MIOX4 OE line compared to WT control (Figure 12A). The *CYP83B1* gene is involved in the indol-and benzyl-glucosinolates biosynthesis which has triptophane and phenylalanine as precursors (Sønderby et al., 2010a). The MYB 76 is transcriptional activator of methionine-derived aliphatic glucosinolates in *Arabidopsis thaliana* (col-0) (Sønderby et al.,2007; Gigolashvili et al., 2008). We found that the level of glucoerucin, a methionine-derived aliphatic glucosinolate, was elevated ∼2.5 fold in *At*MIOX4 OE line compared to WT control (Figure 12B). Interestingly, we found that negative regulator of methionine biosynthesis Arabidopsis MTO 1 responding down 1 (*At*5g53480) was significantly down-regulated in *At*MIOX4 OE line (Supplementary Table 2).

**Figure 12.**
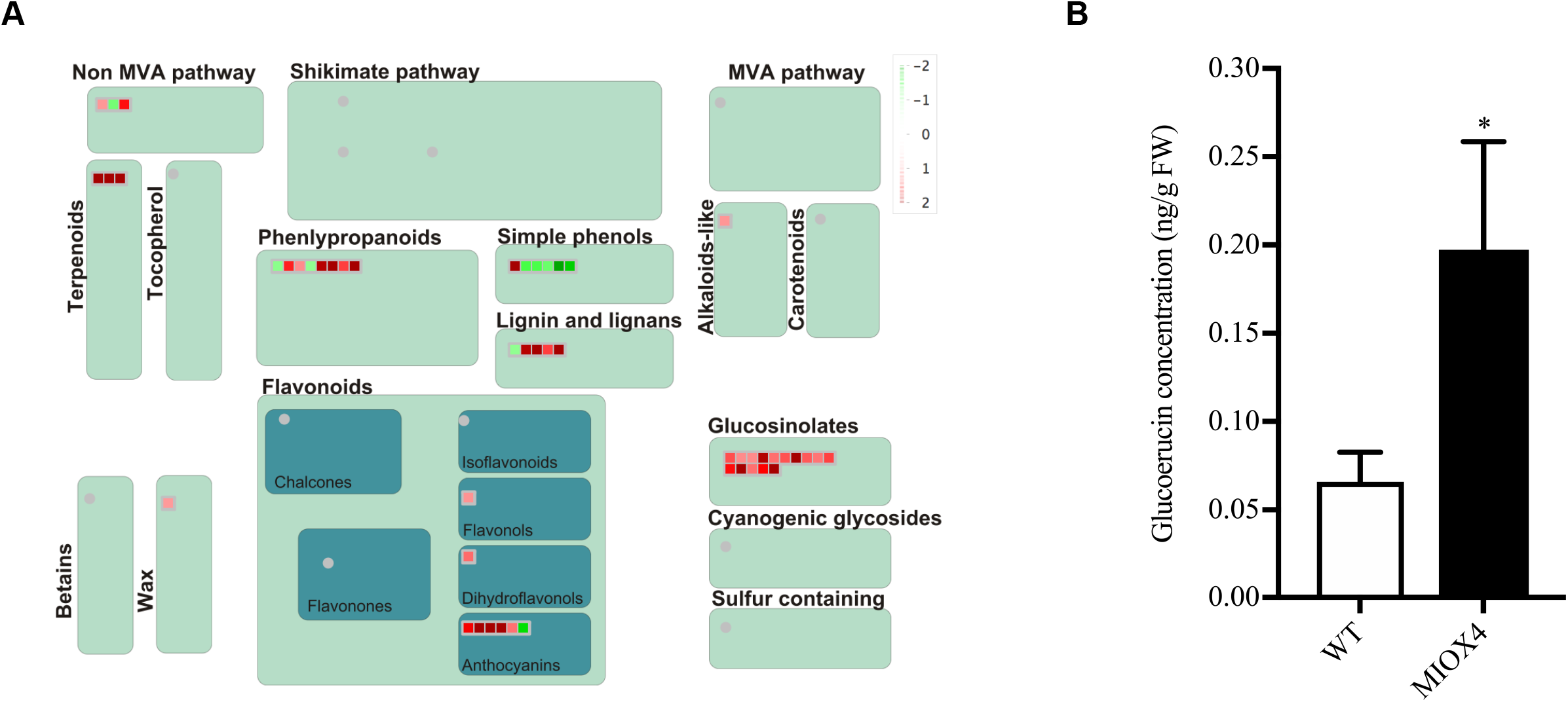
Glucosinolate metabolism was affected by higher ascorbate in plants. A) Glucosinolate metabolism related transcripts were up-regulated in MIOX4 OE line compared to WT line indicated by red square on figure obtained from MapMan v3.5.1R2. B) Glucoerucin concentration was higher in MIOX4 OE line compared to WT line shown by HPLC. Data was mean ± SEM (n=6). * indicated p<0.05 at 0.05 significance level.

### Severity of down-regulation of pathogen resistant genes is compensated by increased SA level

Surprisingly, we found that genes related to the pathogen response as pathogenesis related proteins (*At*2g14610), glycosyl hydrolase superfamily protein (*At*4g16260), peroxidase family proteins (*At*5g51890), germin (*GLP3*: *At*5g51890), leucine rich repeat lipid family protein (*LRR*: *At*5g23400), and disease resistance responsive family protein (*At*1g6580) were down-regulated in the *At*MIOX4 OE line compared to WT controls (Figure 13A). To test whether down-regulation of these genes affected the pathogen defense, we challenged *At*MIOX4 OE (high AsA), *vtc* 1-1 (low AsA), and WT line (moderate AsA) with *Pseudomonas syringae*. As shown in Figure 13B the *At*MIOX4 OE line does not seem to be compromised to defend against *P. syringae* attack, whereas *vtc* 1-1 line were resistant to *P. syringae* challenge compared to WT control when analysed by bacterial count (log cfu/cm^2^) after 3 days of infection (Figure 13B). We evaluated the level of SA in *At*MIOX4 because SA is responsible for protecting the plants against pathogens by provoking the systemic defense response. The foliar level of SA was higher in *At*MIOX4 OE line compared to controls, as shown by LC-MS/MS, without the challenge of biotic stress (Figure 13C). Despite the down-regulated pathogen defense response transcripts, the *At*MIOX4 line was not compromised by the pathogen challenge. This can be due to the presence of high level of SA, which could compensate for the down-regulation of pathogen response genes by expressing lipid transfer proteins related to pathogenesis (*At*3g51590, *At*4g28395, *At*1g66850, *At*5g01870) and *PROTEASE INHIBITOR* (*At*4g22513) genes only in *At*MIOX4 OE line.

**Figure 13.**
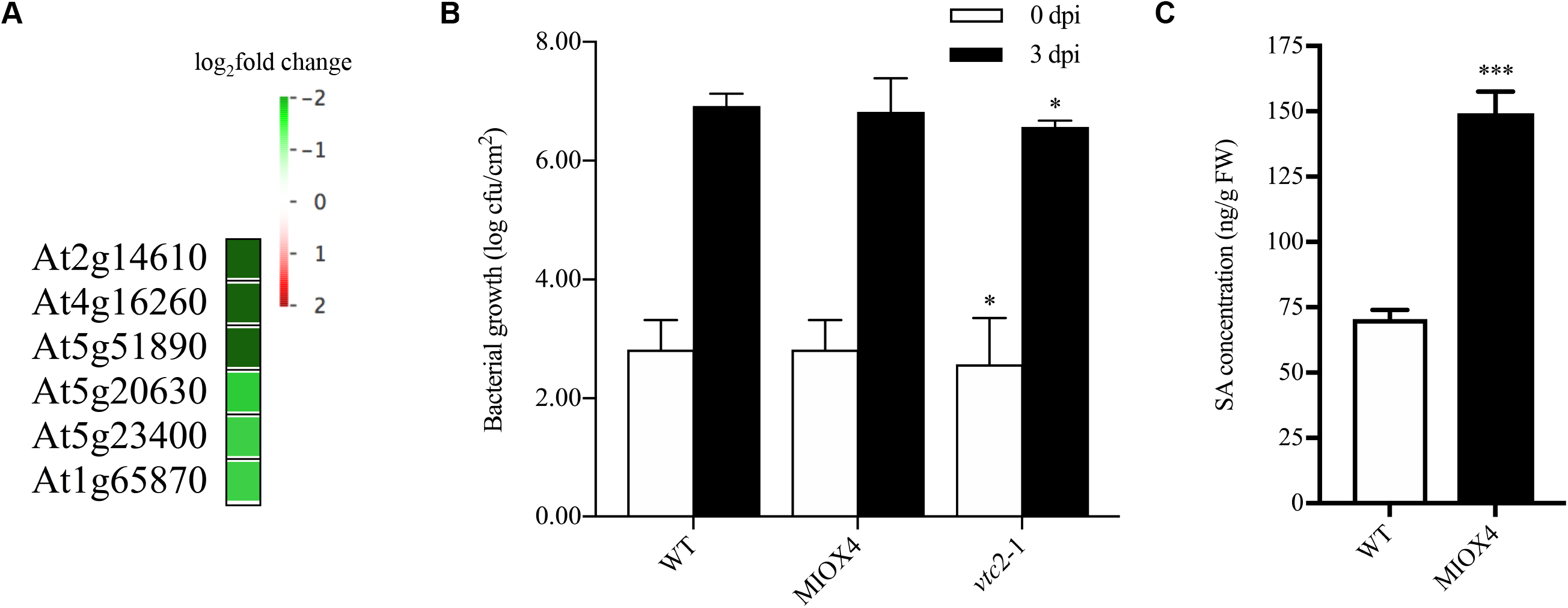
The high ascorbate MIOX4 OE line differentially expressed genes involved in pathogen defense response. A) List of genes that were down-regulated and important for pathogen defense response. B) *Pseudomonas syringae* infection didn’t alter the susceptibility of lines with high ascorbate level. MIOX4 OE and GNL OE were similar to WT level when challenged with *Pseudomonas syringae* whereas low ascorbate lines *vtc*2-1 was resistant to *Pseudomonas syringae* infection. Data was mean ± SD (n=3). C) Defense hormone salicylic acid (SA) was higher in MIOX4 OE line compared to WT line. Data was mean ± SEM (n=5). * indicated p<0.05 and **** indicated p<0.0001 at 0.05 significance level.

## Discussion

Ascorbate is fundamental for proper development and response to biotic and abiotic stress in plants (Conklin et al., 1999; Debolt et al., 2007; Dowdle et al., 2007; Ortiz-Espín et al., 2018; Gallie et al., 2013; Gest et al., 2013; Kerchev et al., 2011; Pastori et al., 2003). The D-mannose/L-galactose pathway (Wheeler et al., 1998) is well characterized compared to other ascorbate routes. Other pathways also called as “alternative routes” are also functional in different plants species, tissues and developmental stage specific manner (Lorence et al., 2004; Yactayo-Chang 2011; Yactayo-Chang et al., 2018). The *MIOX4* transcript is expressed during reproductive stage in leaf stipules and was found predominantly in cytoplasm and in nucleus (Alford et al., 2012; Endres and Tenhaken 2011; Kanter et al., 2005). Over-expression of enzymes involved in the *myo*-inositol pathway leads to increased AsA content in Arabidopsis (Radzio et al., 2003; Lorence et al., 2004; Yactayo-Chang 2011; Yactayo et al., 2018). This has also been confirmed by other laboratories (Tóth et al., 2011). Additionally, a study in a T-DNA insertion mutant of the *vtc*4 enzyme showed that this proteins is involved in *myo-*inositol synthesis resulting in decreased AsA in kiwifruit (*Actinidia deliciosa*) in mutants compared to WT plants (Torabinejad et al., 2018). A recent study reports no conversion of *myo*-inositol into AsA in the AtMIOX4 lines in which we have detected gene silencing (the ones reported in the Lorence et al., 2004 paper) (Kavkova et al., 2018). Because of the gene silencing issue we developed new transgenic lines. The *At*MIOX4 line 21 used in this transcriptomic paper is homozygous, contains a single copy of the transgene, and consistently has shown elevated AsA in more than 10 generations we have grown these plants.

The rosettes of the low ascorbate *A. thaliana vtc*2 mutant are smaller compared to WT due to intracellular structural deformity similar to programmed cell death (Olmos et al., 2006; Pavet et al., 2005). Whereas, plants over-expressing Arabidopsis *myo*-inositol oxygenase (*MIOX4*) (Lorence et al., 2004) have significantly higher (1.5 to 2-fold) ascorbate. Higher AsA results in enhanced biomass, and increased tolerance to abiotic stresses including cold, heat, salt, and pyrene, an environmental pollutant (Lisko et al., 2013). In this study, we report the physiological and molecular mechanisms likely underlying the enhanced biomass and abiotic stresses tolerance phenotype in *At*MIOX4 OE using a comprehensive approach combining transcriptomic analysis, with LC-MS/MS, cell biology and physiological measurements.

We found that an *α-amylase*-like (*At*4g25000) transcript was over-expressed, and resulted in higher intracellular concentration of reducing sugars and glucose in *At*MIOX4 OE line compared to WT using RNA-Seq, RT-qPCR, biochemical assay and enzymatic assay (Figure 4A-D). Glucose acts as a primary carbon and energy source in plants and plays a crucial role in plant growth, development, and physiology by coordinating with phytohormone signals. Enhanced production of glucose leads to manipulation of root architecture for maximum development through interactions with auxin metabolism genes (Li et al., 2007; Moore et al., 2003; Peng et al., 2018). Furthermore, increased concentration of UDP-glucose leads to accumulation of higher biomass in plants and helps in vegetative phase change by repressing the miR156A/ miR156C (Christopher et al., 2017; Yang et al., 2013). Additionally, Glucose induces expression of *MYB* family transcription factors and biosynthetic genes in the aliphatic glucosinolate pathway and in the presence of JA acts synergistically regulating glucosinolates accumulation (Guo et al., 2013: Miao et al., 2016).. Enhanced intracellular glucose accumulation indicates the higher pool of primary carbon and energy source, and signaling molecule for stresses tolerance, which might contribute to higher biomass and abiotic stresses tolerance phenotype in *At*MIOX4 OE line compared to WT.

We have shown that photosynthesis light reaction associated transcripts were up-regulated, and that photosynthetic efficiency and proton motive force were higher in *At*MIOX4 OE line compared to WT (Figure 5A-D). The *At*MIOX4 OE line has higher non-photochemical quenching and energy dependent quenching due to the presence of elevated ascorbate in the lumen. Additionally, in heat stressed leaves of high AsA plants, ascorbate acts as PSII donor by retarding photoinactivation (Tóth et al., 2011).

Photosynthesis light reaction utilizes photosystem II (PSII) to capture the sunlight and splits water molecule into proton, electrons, and oxygen. The free electrons move through the thylakoid membrane and synthesize energy storage molecules. Furthermore, these energy storage molecules are utilized by photosynthesis dark reactions for carbohydrate biosynthesis (Johnson 2016). Higher rate of photosynthesis is associated with higher biomass production and greater yield in plants (Evans 2013; Taylaran 2011). Hence, increase in photosynthesis efficiency in *At*MIOX4 OE provides plant with a higher amount of energy storage molecules and carbohydrates.

The growth and development of plants are regulated by the interaction between growth and defense hormones. We have shown that expression of auxin biosynthesis, auxin transport, auxin-aminoacid conjugate hydrolase, and auxin response signaling factors were up-regulated in *At*MIOX4 OE line compared to the control (Figure 6A-B). The concentration of intracellular active auxin (IAA) and conjugated auxin (IAA-Asp) were higher in *At*MIOX4 OE line compared to WT (Figure 6C). The functional response of elevated auxin in these plants is evidenced by increased primary root length when grown in light conditions, elongated hypocotyl and hypocotyl epidermal cell when grown in dark conditions and both visualization and semi-quantification of auxin levels using confocal microscopy. Auxin concentration was higher in both roots and shoots of *At*MIOX4 OE line harboring the auxin sensor (R2D2) relative to the R2D2 control (Figure 7A-F). Auxin tightly regulates growth and development of plants. Tryptophan dependent auxin biosynthesis transcripts were over-expressed in *At*MIOX4 OE line as nitrilase family gene converts indole-3-acetonitrile to IAA whereas, YUCCA family proteins converts indole-3-pyruvic acid to IAA which is the rate limiting step for tryptophan dependent auxin biosynthesis. Over-expression of both branches of auxin biosynthesis pathway lead to high IAA and IAA-Asp level (Korasick et al., 2013; Zhao et al., 2001). *YUCCA6* over-expression results in high auxin phenotype such as delayed senescence, narrow and downward curled rosette leaves and tolerance to drought stress (Kim et al., 2011; Kim et al., 2013). The over-expression of *YUCCA6, YUCCA1* and external application of auxin derivative delayed senescence and drought stress tolerance in plants due to elevated auxin (Kim et al., 2011; Lee et al., 2011; Mueller-Roeber and Balazadeh 2014). We have found *YUCCA3* over-expression, delayed senescence and abiotic stress tolerance phenotype in the *At*MIOX4 OE line compared to WT control. Additionally, auxin promotes plant organ elongation (hypocotyl, roots and coleoptiles) by enhancing the proton extrusion through the membrane H^+^ ATPase. Auxin lowers apoplastic pH through activation of membrane H^+^ ATPase, which induces cell wall loosening proteins facilitating inward movement of K^+^ and water (Claussen et al., 1997; Philippar et al., 2006; Takahasi et al., 2012; Tyburski et al., 2011). Whereas, cytokinins are involved in cell division and morphogenesis regulation, and abiotic/biotic stresses tolerance (Zalabak et al., 2013). However, constitutive expression of *At*MIOX4 OE did not seem to lead to changes in cytokinin (*cis*-zeatin, *trans*-zeatin and zeatin-riboside) level in *A. thaliana* foliar tissue (Figure 8A-C), which can be observed in both RNA-Seq and LC-MS/MS data.

Analysis of our transcriptomics results and published evidence indicate that abundance of intracellular primary carbon source, increased photosynthetic efficiency and elevated auxin contributes to higher biomass phenotype in *At*MIOX4 OE line compared to WT control. Additionally, delayed senescence in plants was indicated by the presence of high free auxin which is consistent with our results.

### Cell wall metabolism is differentially regulated in AtMIOX4

The cell wall is important to maintain the structural integrity and stresses tolerance of plants. We found that cell wall metabolism genes were differentially modulated in *At*MIOX4 OE line compared to WT control (Figure 11A; Supplementary Table 3). Lignin maintains the structure of the secondary cell wall. We found that laccase biosynthesis genes (*LAC11* and *LAC17*) were down-regulated in *At*MIOX4 OE line compared to WT control. Knockout of laccase biosynthesis (*LAC1* and *LAC17*) genes in *A. thaliana* results in reduced plant growth and vascular development, decreased root diameter due to reduced lignification and increased saccharification (Berthet 2011; Zhao et al., 2013) which contrasts with our results. Auxin decreases the expression of *AGP3* whereas *AGP19* is involved in plant growth, cell division and expansion, leaf development and reproduction (Goda et al., 2004; Yang et al., 2007). Interestingly, *A. thaliana* arabinogalactan protein (*At*AGP17) over-expression results in resistance to infection by *Agrobacterium* (Gaspar et al., 2004). The expression of the gene *longofolia* is involved in polar cell expansion by turgor driven approach by controlling *XTH17* and *XTH24* expression (Lee et al., 2017). In contrast, *CSLA/CSLA-*like gene family members that are involved in mannan and xyloglucan backbones biosynthesis (Liepman and Cavalier 2012) were up-regulated in *At*MIOX4 OE line compared to WT counterpart.

### Abiotic stress tolerance of high ascorbate line were independent of increased abscisic acid level

Prior studies have shown that, low ascorbate *A. thaliana* lines were sensitive to abiotic stresses such as heat, cold, light and oxidative stress (Conklin et al., 2013; Pavet et al., 2005). On the other hand, plants with elevated ascorbate were tolerant to abiotic stresses as heat, cold, drought, salt and oxidative stress (Eltayeb et al., 2012; Eltayeb et al., 2006; Lisko et al., 2013; Tóth et al., 2011; Yactayo-Chang et al., 2018). We found that heat stress response genes (*HSP17.8-CI; HSP17.4-CIII*), cold stress response genes (*COR15A*), and drought stress response genes (*ATDI21, NAC019*) were up-regulated in the *At*MIOX4 OE line compared to WT (Figure 9A-B; Supplementary Figure 3). Whereas, intracellular concentration of ABA was similar in *At*MIOX4 OE line and WT (Figure 9C). Over-expression of class I and class II HSPs are important for basal thermotolerance by interacting with translation factor (eEF1B), and knockdown of class II HSPs by RNAi result in severe heat sensitivity (McLoughlin et al., 2016). Moreover, over-expression of *HSP 17.8* aggregates the *A. thaliana* citrate synthase enzyme and protect it from heat at 43°C (Liu et al., 2005). Additionally, *HSP 17.8* over-expression results in freezing tolerance in maize (Shou et al., 2004). Similarly, over-expression of *HSP 17.4-CII* results in heat stress tolerance in tomato through co-regulation of heat stress induced heat shock transcription factor A2 (HsfA2) (Port et al., 2004). Constitutive expression of *Cor15A* enhances freezing tolerance of protoplast and chloroplast in *A. thaliana.* This is due to decrease incidence of lamellar to hexagonal II phase transitions which occur in the close proximity of chloroplast membrane and plasma membrane due to dehydration caused by freezing (Artus et al., 1996; Steponkus et al., 1998; Thalhammer et al., 2014). Interestingly, over-expression of *NAC* family transcription factors (*ATAF, CUC* and *NAM*) in rice enhances drought resistance and salt tolerance by trans-activating activity (Hu et al., 2006). In addition to this, over-expression of *ANAC019* increases heat tolerance and drought tolerance in *A. thaliana* (Tran et al., 2004; Guan et al., 2004).

Peroxisomal ascorbate peroxidase 5 (*pAPX5*) is involved in scavenging ROS, which is down-regulated in the *At*MIOX4 OE line compared to WT (Figure 9A-B). High amount of ROS are produced in peroxisomes due to peroxisomal specific enzymes as acyl coA oxidase, glycolate oxidase, sulphite oxidase, superoxide dismutase, sarcosine oxidase, and polyamine oxidase in different plant species (Arent et al., 2008; Douce and Neuberger 1999; Goyer et al., 2004; Hansch et al., 2006; del Rio et al., 2006; Kamada et al., 2008; Planas et al., 2013). Reactive oxygen species produced in peroxisomes are scavenged by *pAPX* or catalase (CAT). Spinach *pAPX* have high affinity for H_2_O_2_ compared to catalase (Ishikawa et al., 1998). Reduction in expression of *pAPX* results in higher level of H_2_O_2._ Enhanced level of H_2_O_2_ in peroxisomes results in up-regulated expression of genes involved in oxidative stress response as small heat shock family proteins, DNAJ heat shock family protein, aromatic alcohol: NADPH reductase, and bHLH transcription factor. It was previously observed that silencing of *pAPX* results in oxidative stress tolerance in rice (del Rio et al., 2006; Sousa et al., 2015).

Based on a combination of the results shown here and published evidence, we propose a model were elevated AsA in *A. thaliana* reduces the expression of *pAPX* resulting in high level of peroxisomal ROS. High ROS in peroxisomes may act as signaling molecules and elevate the expression of abiotic stresses tolerance genes as heat response genes (*HSP17.8-CI; HSP17.4-CIII*), DNAJ heat shock protein family, bHLH transcription factor and aromatic alcohol: NADPH reductase. This will result in plants tolerant to oxidative stress. In addition to this, cold stress response gene (*COR15A*), drought stress response genes (*ATDI21, NAC019*) that were up-regulated in *At*MIOX4 OE line could lead to cold and drought stress tolerance.

### High AsA plants seem ready to defend themselves from stresses

Genes related to specialized metabolism were differentially regulated in *At*MIOX4 OE line compared to WT control. Phenolics biosynthesis gene o-methyl transferase family 2 protein was up-regulated. Phenylpropanoid biosynthesis genes (*TSM1, CAD-B2*), flavonoid biosynthesis (*MYB75*), the *JAO2, JAO3, JAO4, AACT1* genes and terpene biosynthesis genes (*GGPS6, LAS1, TPS10*) were up-regulated in the high AsA line compared to controls (Figure 11A). Over-expression of *MYB75* results in accumulation of anthocyanin and protects *A. thaliana* from high light intensity (Gonzalez et al., 2008; Li et al., 2016). Glucosinolates-myrosinase system genes such as flavin monoxygenase glucosinolate s-oxygenase 2, alkenyl hydroxalkyl producing 2, MYB domain protein 76, myrosinase binding protein 1, and epithiospecifier protein were up-regulated in *At*MIOX4 OE line compared to WT (Figure 12A). *MYB76* transcription factor positively regulates the biosynthesis of methionine-derived aliphatic glucosinolates (Sønderby et al, 2007; Gigolashvili et al., 2008). *MYB76* is involved in synthesis of methylsulfinylalkyl glucosinolates from methylthioalkyl glucosinolates catalyzed by FMO_GSOX_S enzyme family (Hansen et al. 2007; Li et al., 2008). Epithiospecifier protein is involved in nitrile formation upon glucosinolates hydrolysis and making Arabidopsis less appealing to specialist herbivores (Burow et al., 2010; Lambrix et al., 2001). Methionine-derived glucosinolates acts as a chemoprotective compounds in plant defense against biotic stress (Gigolashvili et al., 2007). Increased levels of Glucoerucin observed in *At*MIOX4 OE might indicate that high AsA plants could be less susceptible to herbivore attack. Furthermore, Glucoerucin, when converted into erucin by myrosinase enzyme, acts as an antioxidant by decomposing hydroperoxides and hydrogenperoxides. Erucin oxidizes into glucoraphanin which is most effective chemoprotective for cancer. Eruin protects from cancer by multiple mechanisms as modulation of phase I enzymes, inducing phase III enzymes, up-regulating phase III detoxification system, and modulation cell proliferation (Barillari et al. 2005; Dinkova-Kostova et al. 2017; Fimognari et al. 2017; Melchini and Traka 2010). Further analysis on other glucosinolates present in *At*MIOX4 OE line is now in progress and will help us to understand the effect of glucosinolates in this plant.

Jasmonic acid biosynthesis gene and JA-responsive genes were up-regulated in *At*MIOX4 OE line compared to WT. JA biosynthesis intermediate metabolites (OPDA) were similar but JA and JA-Ile level were significantly higher in *At*MIOX4 OE line compared to controls (Figure 11A-E). Over-expression of *OPR3* results in increased level of JA and JA-Ile in *A. thaliana* and plants become resistance to insects and fungi (Suza and Staswick 2008) which is similar to our results. Interestingly, JA is also involved in plant development, growth, reproduction, and protects plants against abiotic stresses as salinity, drought, and UV-radiation (Kerchev et al., 2011; Wasterneck and Hause 2013). Salicylic acid metabolism genes and SA level were up-regulated in *At*MIOX4 OE line compared to WT control. In contrast, pathogenesis related proteins were down-regulated in *At*MIOX4 OE line compared to WT control (Figure 11A, 13A, 13C). The expression of pathogenesis related proteins was significantly down-regulated in *At*MIOX4 OE line. Surprisingly, the response of *At*MIOX4 OE line was similar to WT control when challenged with *P. syringae.* This may be due to exclusive expression of pathogen response genes in *At*MIOX4 line (Figure 3A; Table 2). Salicylate protects the plants from biotrophic and hemibiotrophic pathogens by interacting with *NPR1* (Palmer et al., 2017). Salicylic acid not only protect plants against biotic stress, it is also involved in growth, development of plants and increases tolerance against abiotic stresses as cold, heat, drought, metal by increasing antioxidant capacity (Bartoli et al., 2013; Gimenez et al., 2017; Horváth et al., 2015). Furthermore, ethylene-responsive element binding factor 13 induces the over-expression of RAP transcriptions factor. Over-expression of RAP2.6 resulted in increased resistance to nematode (*Heterodera schachttii*) in *A. thaliana* by enhanced callose deposition (Ali et al., 2013).

### Crosstalk between high AsA, SA and JA

Glutathione reductase 480 (*GRX480*) and *DHAR1* which is involved in recycling of reduced ascorbate were both up-regulated while peroxisomal p*APX5* was down-regulated in *At*MIOX4 OE line compared to WT control (Figure 9A-B). It was reported that, under high light stress, elevated H_2_O_2_ is associated with up-regulation of small heat shock proteins and transcription factors related to anthocyanin biosynthesis (Vanderauwera et al., 2005). Similarly, anthocyanins protect plants against photo-protection and photo-oxidative damage. Mutation of thylakoid *APX* results in elevated H_2_O_2_ under high light stress which increases the expression of genes involved in anthocyanin biosynthesis regulation (Maruta et al., 2014). We found that the positive regulator of anthocyanin biosynthesis *MYB75* and the anthocyanin biosynthesis 2OG-Fe(II) oxygenase family proteins were up-regulated in *At*MIOX4 OE line compared to control (Supplementary Table 3). These results suggest that high AsA *A. thaliana* are ready to protect themselves from photo-oxidative damage by regulating anthocyanin biosynthesis.

Salicylic acid negatively regulates the JA biosynthesis genes and decreases expression of JA-responsive genes (*PDF1.2* and *VSP2*) (Chehab et al., 2011; Vlot et al., 2008; Leon-Reyes et al., 2010). We found that JA and SA were both elevated and JA-responsive defensin proteins were highly expressed in *At*MIOX4 OE line compared to control. *GRX480* was inducible by SA and suppresses the JA responsive defensin (*PDF1.2*) transcription in a *NPR1* dependent manner. *GRX48*0 interacts with *TGA2* transcription factor and suppresses the ethylene-response factor 59 *(ERF59)* resulting in suppression of JA response defensin (Ndamukong et al., 2007). Published literature indicates that higher levels of endogenous ethylene diminish the SA and JA antagonism (Leon-Reyes et al., 2009; Leon-Reyes et al., 2010). Ethylene response factors as *RAP2.6, RAP2.6L, RAP 2.10* were up-regulated in *At*MIOX4 OE line compared to WT control (Figure 11A). The over-expression of the ethylene response factors might contribute to diminish the SA and JA antagonism, hence increased level of both SA and JA in high AsA *A. thaliana*.

### Conclusions and future perspectives

We have here demonstrated that high AsA *At*MIOX4 OE contain increased intracellular glucose, and enhanced photosynthetic efficiency. We also found that elevated *in vivo* auxin levels contributed to increased cell elongation and delayed senescence. The enhanced glucose, auxin and photosynthetic efficiency are likely explanations for the enhanced biomass phenotype of the high AsA line. Furthermore, the up-regulated expression of signature genes and transcription factors for heat stress, cold stress, drought stress and salt stress tolerance over-expression likely lead to abiotic stresses tolerance in this high AsA *At*MIOX4 OE line. The fact that abscisic acid was similar in *At*MIOX4 and controls indicates that abiotic stresses tolerance in this line is most likely independent of up-regulation of ABA. Interestingly, biotic stress response genes related to JA, SA, specialized metabolism and glucosinolates biosynthesis and metabolism genes were up-regulated in the high AsA line. The defense hormones JA, JA-Ile and SA were elevated in the high AsA line compared to its counterparts. These defense hormones and elevated expression of biotic stress response signature genes might confer the tolerant/resistant phenotype of *At*MIOX4 in response to biotic stresses. Furthermore, the up-regulated expression of ethylene response factors in the high AsA line strongly suggests the presence of elevated levels of ethylene. Elevated ethylene might abolish the SA and JA antagonism in high AsA line. These results not only shed light into the molecular mechanisms underlying the enhanced biomass and abiotic stress tolerance phenotype of the high AsA line but also point to a conserved module of biotic stress response genes. Future studies are necessary to test the ability of high AsA lines to respond when challenged with herbivores. Next, steps should also include the expression of *At*MIOX4 in crops of commercial value to fully exploit the potential biotechnological value of this knowledge.

## Supporting information

Suppementary Table 2

## Acknowledgments

We thank Dr. Fabricio Medina-Bolivar (A-State) for providing access to the HPLC, Dr. Sophie Alvarez (UNL) for technical assistance in LC-MS/MS measurements, Dr. Jian Hua (Cornell) for assistance with pathogens assays, Krishna Deo Sharma (A-State) for assistance in microscopy, Zachary Campbell (A-State) for technical assistance, and Kimberly Lee (A-State) for plant care. We thank Dr. Fiona Goggin (UAF) for insightful suggestions to the project. NN thanks the Molecular Biosciences PhD Program for providing stipend support. We also thank the Arkansas Biosciences Institute (ABI, A-State) for providing facilities to conduct experiments. This research was funded by the Arkansas Center for Plant Powered Production (Arkansas EPSCoR Track 1, fund EPS 0701890), the Plant Imaging Consortium (PIC, AR EPSCoR Track 2, fund IIA 1430427) funded by the National Science Foundation and a grant to AL from the Arkansas Research Alliance.

## Author Contributions

NN performed the experiments, prepared figures, and wrote initial draft of the manuscript, JPYC generated the MIOX4 homozygous line; JPYC and LMAG prepared RNA samples for sequencing, MEGR contributed to HPLC analysis, KMJ and MAV analyzed the RNA-Seq data, AL conceived the experiments, secured funding, and wrote final draft of the manuscript.

## Supplementary materials

**Supplementary Figure 1.** The differential expression of genes involved in multiple L-AsA biosynthesis pathways.

**Supplementary Figure 2**. UV-spectra and chromatogram of glucoerucin at 229 nm.

**Supplementary Figure 3.** Genevestigator analysis of top 200 up-regulated transcripts (log_2_fold > 1.75) were compared with signature expression transcripts.

**Supplementary Figure 4.** Down-regulation of cell wall metabolism transcripts.

**Supplementary Figure 5.** ATTEDII analysis of up-regulated transcripts.

**Supplementary Table 1.** Up-regulated transcripts lists in *At*MIOX4 OE line.

**Supplementary Table 2.** Down-regulated transcripts lists in *At*MIOX4 OE line.

**Supplementary Table 3.** MapMan transcripts and functions.

**Supplementary Table 4.** Up-regulated GO-terms list.

**Supplementary Table 5.** Down-regulated GO-terms list.

**Supplementary Table 6.** Primers used in this study.

