## Supplementary Figures for "Mechanisms Underlying the Enhanced Biomass and Abiotic Stress Tolerance Phenotypes of an Arabidopsis MIOX Over-expresser"

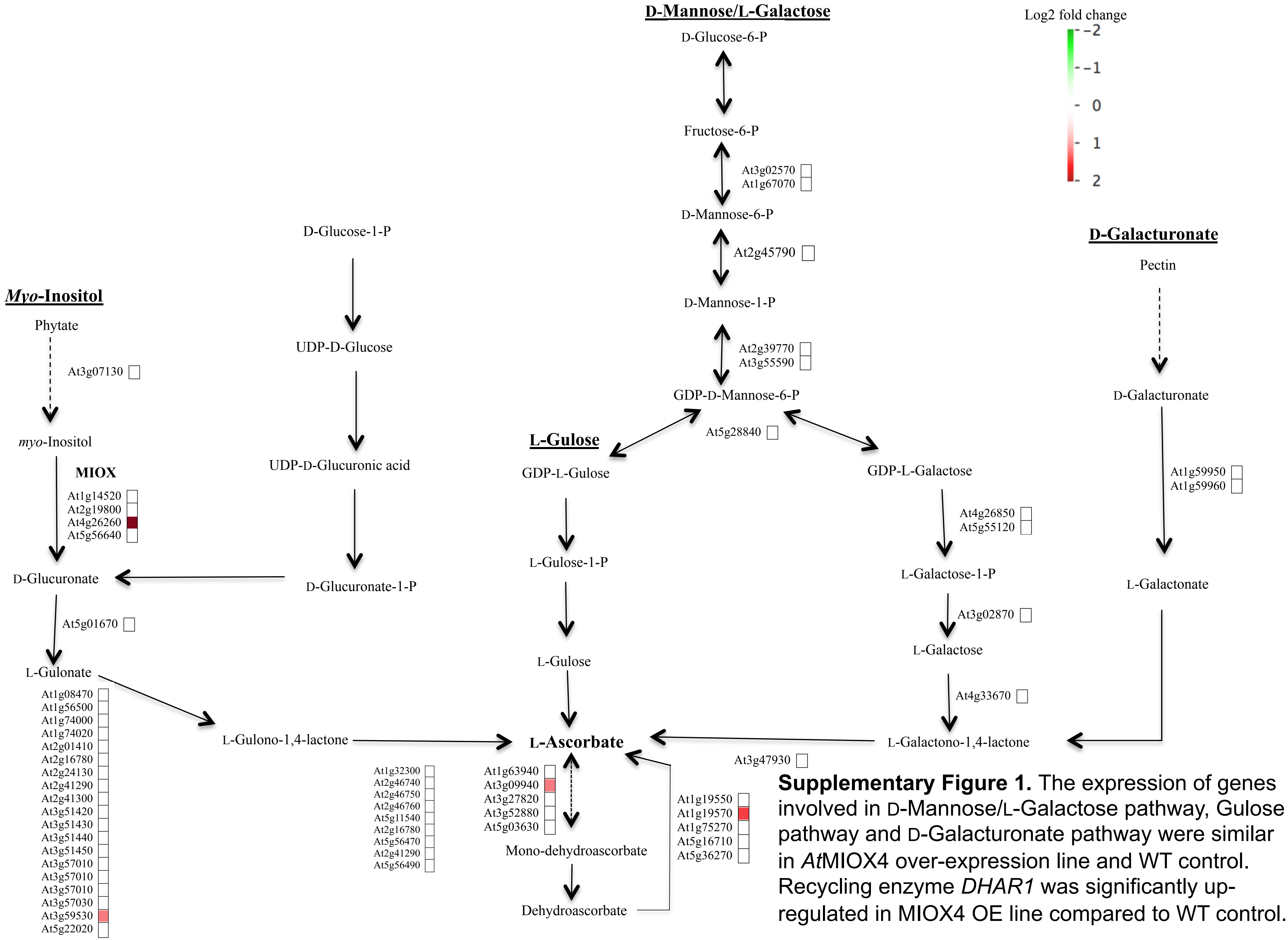


**Supplementary Figure 1.** The expression of genes involved in D-Mannose/L-Galactose pathway, Gulose pathway and D-Galacturonate pathway were similar in *At*MIOX4 over-expression line and WT control. Recycling enzyme *DHAR1* was significantly up-regulated in MIOX4 OE line compared to WT control.

**A** **B**

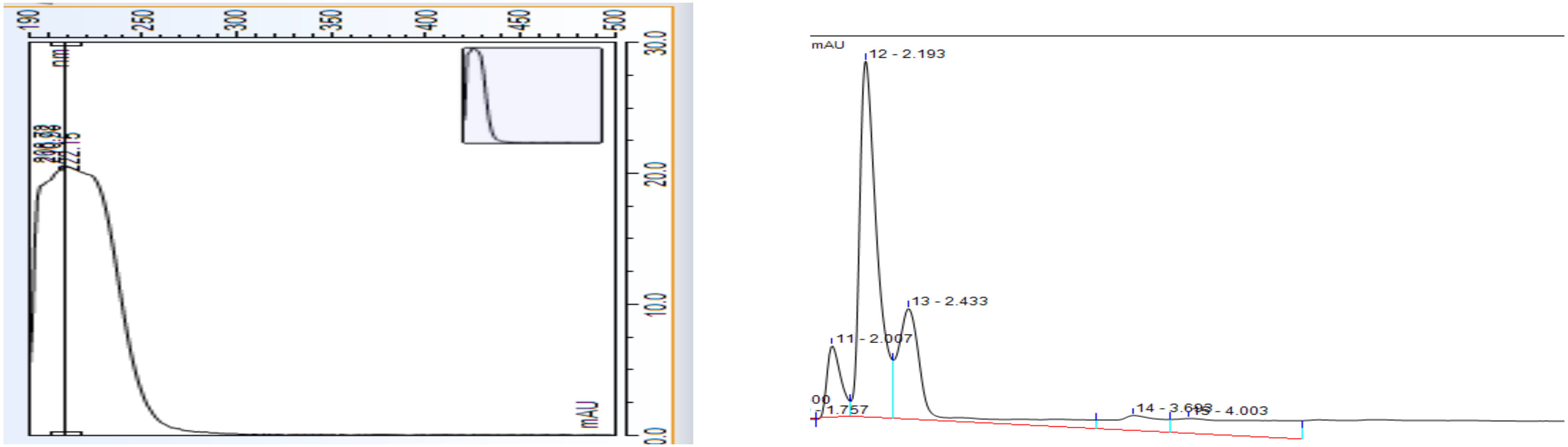


**Supplementary Figure 2**. A) UV-spectra of glucoerucin at 229nm B) Chromatogram of glucoerucin detected at 229nm.

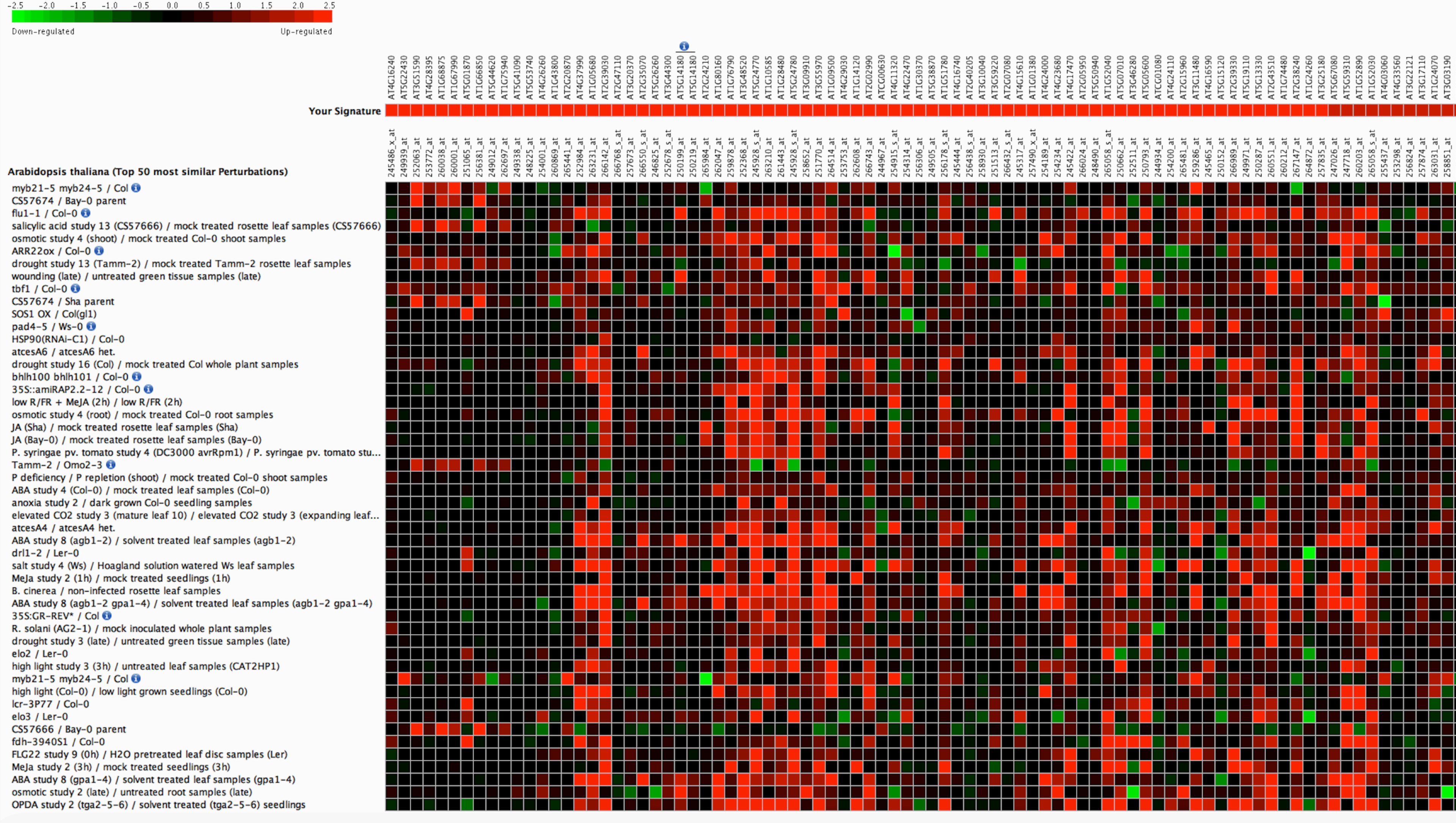


**Supplementary Figure 3.** GeneVestigator analysis of top 200 up-regulated transcripts ($log_2$fold > 1.75) were compared with signature expression transcripts from 50 abiotic and biotic stress perturbations. *ATCAD8* , *GLYI7* and *NAC019* were also highly expressed in most of the abiotic stress (heat, salt, drought cold) tolerance experiments.

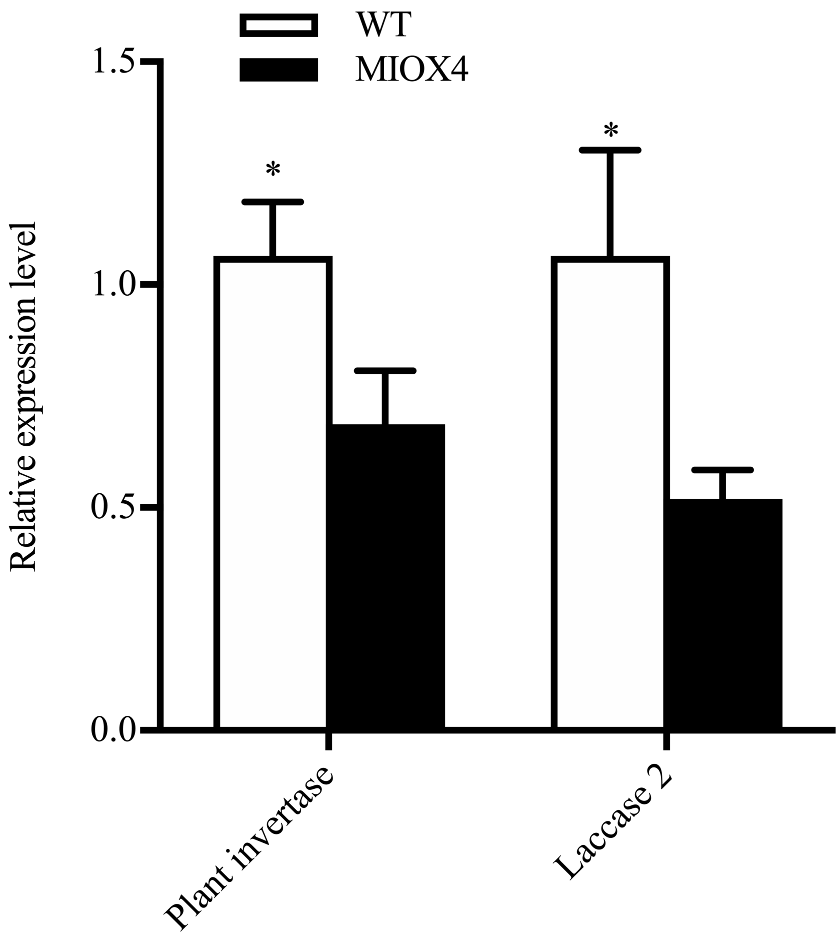


**Supplementary Figure 4**. Down-regulation of cell wall metabolism transcripts. Validation of down-regulation of plant invertase/ pectinmethylestarase superfamily protein and Laccase 2 in *At*MIOX4 OE line using RT-qPCR.  * indicated $p<0.05$ at 0.05 significance level. Data was mean ± SEM (n=3).

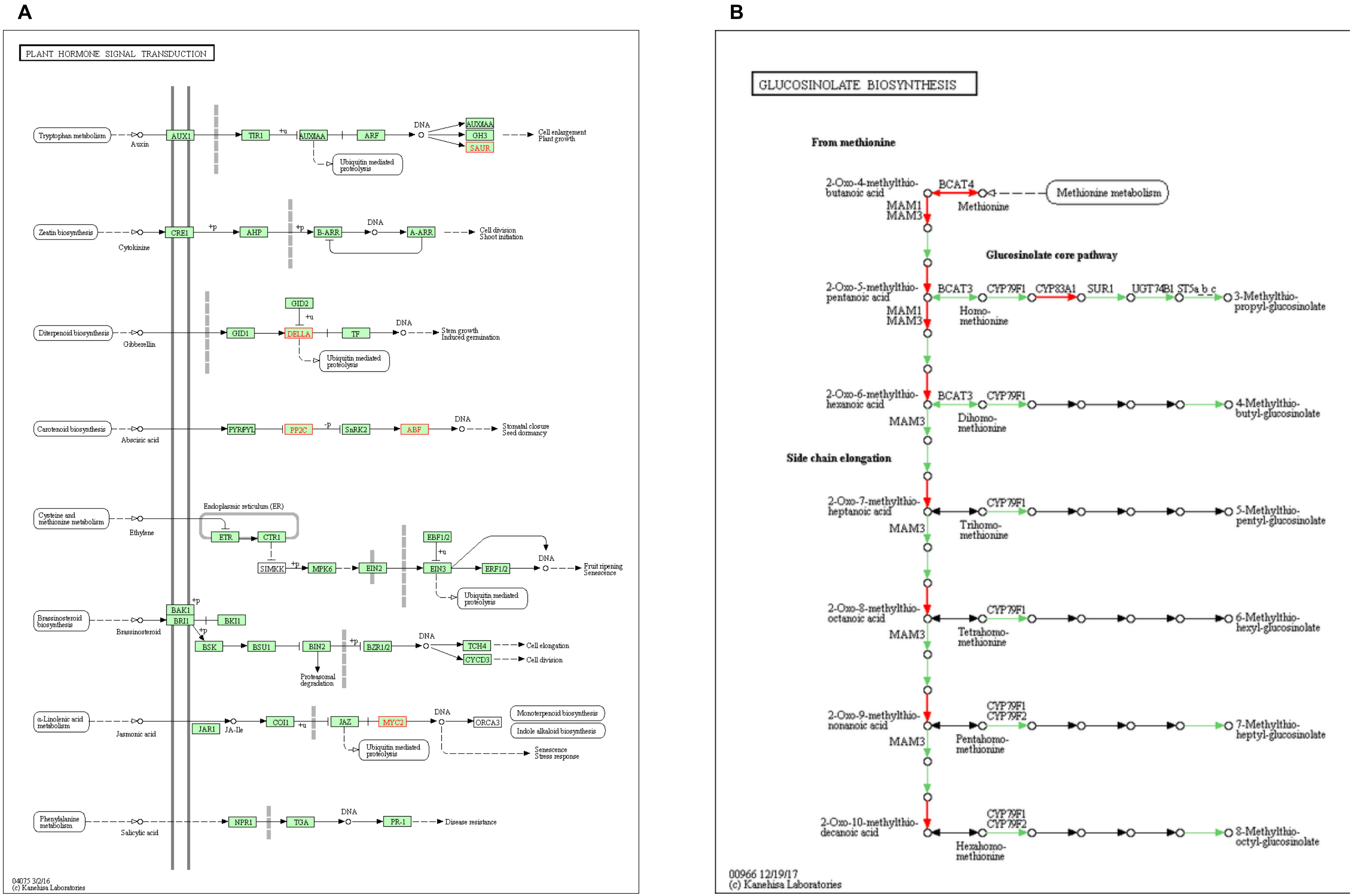


**Supplementary Figure 5**. ATTEDII analysis of up-regulated transcripts. A) Mapping of up-regulated transcripts in plant hormone signaling pathway, B) Mapping of up-regulated transcripts in glucosinolates biosynthesis pathway.
